# Targeting EIF4A triggers an interferon response to synergize with chemotherapy and suppress triple-negative breast cancer

**DOI:** 10.1101/2023.09.28.559973

**Authors:** Na Zhao, Elena B. Kabotyanski, Alexander B. Saltzman, Anna Malovannaya, Xueying Yuan, Lucas C. Reineke, Nadia Lieu, Yang Gao, Diego A Pedroza, Sebastian J Calderon, Alex J Smith, Clark Hamor, Kazem Safari, Sara Savage, Bing Zhang, Jianling Zhou, Luisa M. Solis, Susan G. Hilsenbeck, Cheng Fan, Charles M. Perou, Jeffrey M. Rosen

**Author notes:** Correspondence to Jeffrey Rosen.

## Abstract

Protein synthesis is frequently dysregulated in cancer and selective inhibition of mRNA translation represents an attractive cancer therapy. Here, we show that therapeutically targeting the RNA helicase eIF4A by Zotatifin, the first-in-class eIF4A inhibitor, exerts pleiotropic effects on both tumor cells and the tumor immune microenvironment in a diverse cohort of syngeneic triple-negative breast cancer (TNBC) mouse models. Zotatifin not only suppresses tumor cell proliferation but also directly repolarizes macrophages towards an M1-like phenotype and inhibits neutrophil infiltration, which sensitizes tumors to immune checkpoint blockade.

Mechanistic studies revealed that Zotatifin reprograms the tumor translational landscape, inhibits the translation of *Sox*4 and *Fgfr1*, and induces an interferon response uniformly across models. The induction of an interferon response is partially due to the inhibition of *Sox4* translation by Zotatifin. A similar induction of interferon-stimulated genes was observed in breast cancer patient biopsies following Zotatifin treatment. Surprisingly, Zotatifin significantly synergizes with carboplatin to trigger DNA damage and an even heightened interferon response resulting in T cell-dependent tumor suppression. These studies identified a vulnerability of eIF4A in TNBC, potential pharmacodynamic biomarkers for Zotatifin, and provide a rationale for new combination regimens comprising Zotatifin and chemotherapy or immunotherapy as treatments for TNBC.

**One Sentence Summary:** Targeting EIF4A sensitizes TNBC to immune therapy and chemotherapy by suppressing Sox4, inducing an interferon response, and reprograming the tumor immune microenvironment.

## Introduction

Triple-negative breast cancer (TNBC) is a heterogeneous group of breast cancers defined by the absence of estrogen receptor (ER), progesterone receptor (PR), or HER2, and is minimally classified into four genomic subtypes (1). TNBCs patients have pathological complete response (pCR) rates of 30–53% when treated with a neoadjuvant anthracycline/taxane-containing regimen (2) and recently in a subset of patients pCR rates were improved following treatment with immune checkpoint blockade (ICB) (3). Recent improvements have also been made in treating metastatic TNBC with an antibody-drug conjugate (4) or ICB in combination with chemotherapy in PD-L1+ TNBC (5, 6). However, there is a critical need to identify therapeutic vulnerabilities and treatments that can potentiate the response to chemotherapy and immunotherapy.

One key target for cancer therapy is protein synthesis, which is frequently dysregulated in cancer. Nearly all critical oncogenic signaling pathways ultimately rewire the translational machinery to support tumorigenesis (7). Of all steps in protein synthesis, translation initiation is the rate-limiting step and is subject to extensive regulation (8). Oncogenic signaling pathways promote translation initiation mainly through stimulation of the eukaryotic translation initiation factor (eIF) 4F complex. The eIF4F complex comprises three components, of which eIF4A is an RNA helicase that catalyzes the unwinding of secondary structure in the 5′-untranslated region (5′-UTR) of mRNAs to facilitate translation initiation (9). Multiple tumor-promoting genes that contain structured 5’-UTRs require enhanced eIF4A activity for translation (10–13). The natural compound Rocaglamide A (RocA) and its derivatives (Rocaglates) can target the bimolecular cavity formed by eIF4A and polypurine RNA (14). This interaction of Rocaglate clamps eIF4A on mRNAs that contain polypurine motifs in their 5’-UTR and results in sequence-selective translation repression and translatome remodeling by blocking scanning of the pre-initiation complex and other mechanisms (15–17). This is contrary to Hippuristanol, an initiation inhibitor that binds to the C-terminus of eIF4A and blocks RNA binding in a sequence-independent manner (18, 19). Several studies have reported a dependency on eIF4A in different cancers, suggesting the therapeutic potential of Rocaglates (12, 13, 20–22). However, these natural compounds do not possess good drug properties and their effects on the tumor immune microenvironment are not well-defined.

Zotatifin (eFT226) is a chemically designed Rocaglate derivative with improved drug-like properties and is the first-in-class eIF4A inhibitor (23). Zotatifin is currently undergoing a Phase I/II clinical trial in KRAS-mutant tumors and ER+ breast cancers (NCT04092673). However, TNBC patients are not enrolled in the trial due to a lack of preclinical studies, and pharmacodynamic biomarkers for Zotatifin remain to be identified. Zotatifin has been shown previously to inhibit tumor growth in several immunocompromised mouse models (20). However, since the immune system plays a crucial role in both tumor development and treatment response, it is crucial to examine Zotatifin in immune-competent preclinical models. Moreover, studies on the interaction of Zotatifin with chemotherapy are important because chemotherapy remains the primary systemic treatment option for patients with TNBC and many other solid cancers. We previously developed multiple novel syngeneic TNBC genetically engineered mouse (GEM) models across different intrinsic molecular subtypes, which have been characterized both genetically and with respect to their immune microenvironments (24–30). Here we tested the therapeutic efficacy of Zotatifin both as a monotherapy and in combination with immunotherapy or chemotherapy in these GEM models. We found that Zotatifin exerts pleiotropic effects on both tumor cells and the immune microenvironment and synergizes with carboplatin in mounting an interferon response resulting in a T cell-dependent suppression of tumor growth.

## Results

### Zotatifin monotherapy inhibits tumor growth in a cohort of syngeneic *Trp53*-null mammary tumor models

As *TP53* is the most frequently altered gene in TNBC (31), syngeneic TNBC GEM models were generated previously by *in situ* transplantation of donor mammary epithelium from *Trp53*-null BALB/c mice into wild-type recipient hosts (**Fig. 1A**). This resulted in the derivation of heterogeneous *Trp53*-null transplantable mammary tumors. Detailed genomic characterization has revealed that these tumors are representative of the different intrinsic molecular subtypes of human breast cancer, including the basal-like, luminal-like and claudin-low subtypes (24–27). Besides the apparent differences in tumor histology, these tumor models also exhibited variable infiltration of myeloid cells including macrophages and neutrophils as illustrated by the immunostaining of F4/80 (macrophage marker) and S100A8 (neutrophil marker) (32), respectively (**Fig. 1B**). Tumors of the Claudin-low subtype (T12 and 2151R) were more mesenchymal and highly infiltrated with macrophages (**Fig. 1B**). This correlation of mesenchymal features with macrophage infiltration has also been observed in human TNBC (33). We first determined the effect of Zotatifin monotherapy in six GEM models across three subtypes. To minimize intertumoral variation between mice, we transplanted freshly dissociated tumor cells instead of tumor chunks into the mammary fat pad of BALB/c mice. When tumors reached approximately 100 mm^3^ in volume, we randomized the mice and started treatment with either vehicle or Zotatifin at 1 mg/kg every three days (**Fig. 1C**). Most of these models are so aggressive that they rapidly reach the ethical endpoint within one to two weeks after randomization, but Zotatifin treatment slowed tumor growth without obvious bodyweight loss except for those caused by tumor weight reduction in 2225L-LM2 (**Fig. 1D; Supplementary Fig. S1A and B**). Interestingly, in 2151R, vehicle-treated mice started to lose bodyweight starting at day 9, possibly due to cachexia induced by enlarged tumors, but mice in the Zotatifin treatment group were not affected (**Supplementary Fig. S1B**). Besides GEM models, we also tested 4T1 and E0771, two TNBC cell line models. Whereas Zotatifin treatment reduced the 4T1 tumor volume, E0771 was completely resistant (**Supplementary Fig. S1C**). Neither model showed bodyweight loss (**Supplementary Fig. S1D**). These data suggest that Zotatifin is an effective and well tolerated therapy for the majority of TNBC models.

**Figure 1.**
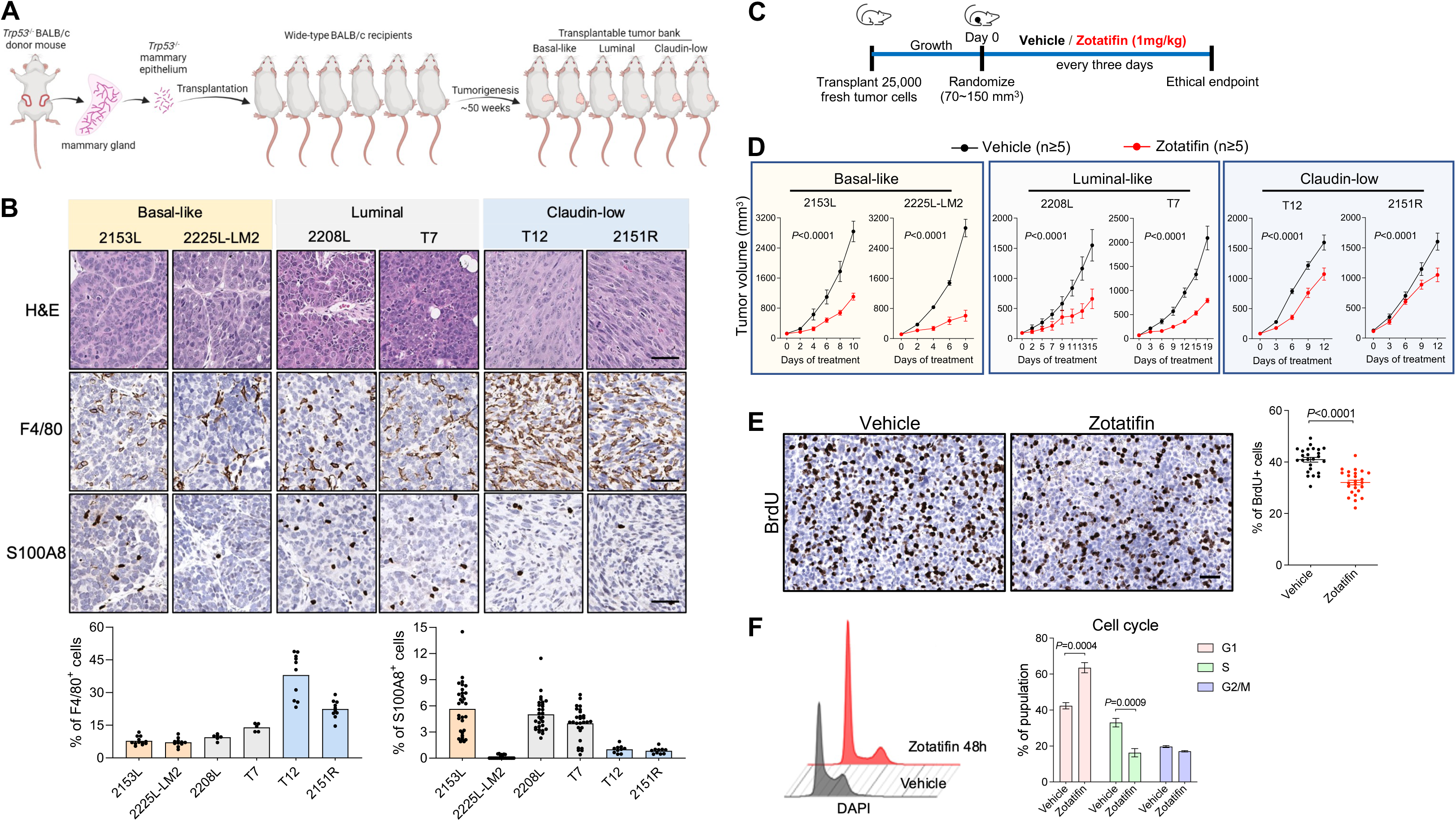
Zotatifin monotherapy inhibits tumor growth in a cohort of *Trp53*-null preclinical models. **A,** Scheme depicting the generation of syngeneic *Trp53*-null preclinical models. Donor mammary epithelium from *Trp53*-null BALB/c mice was transplanted *in situ* into cleared mammary fat pad of wild-type recipient hosts, resulting in the derivation of heterogeneous *Trp53*-null transplantable mammary tumors representative of the different intrinsic molecular subtypes of human breast cancer. **B,** Top, representative images of H&E staining and IHC staining of F4/80 and S100A8 in *Trp53*-null models used in this study. Scale bar, 50 μm. Bottom, quantification of IHC staining. Three to six representative 20X images were analyzed for each tumor. **C,** Schematic of animal treatment. Freshly dissociated tumor cells were injected into the 4^th^ mammary fat pad of BALB/c mice. Mice were randomized and treatment was initiated when tumors reach 70∼150 mm^3^ volume. Mice were treated with either vehicle or Zotatifin every three days until ethical endpoint. **D**, Tumor growth curves of BALB/c mice treated with either vehicle or Zotatifin. n=6 biological replicates for 2225L-LM2 and n=5 for all other models in each treatment arm. Data are presented as mean ± SEM and analyzed using two-way ANOVA with Bonferroni’s multiple comparison test. **E,** Left, representative images of IHC staining of BrdU in ethical endpoint 2153L tumor tissues. Scale bar, 50 μm. Right, quantification of IHC staining. Five representative 20X images were analyzed for each tumor. n=5 biological replicates. Data are presented as mean ± SEM and analyzed using two-tailed unpaired Student’s *t*-test. **F,** Left, representative images of cell cycle distribution of 2153L cells that were cultured in complete medium and treated with vehicle or 40 nM Zotatifin for 48 hrs. Right, quantification of the cell cycle phases from three independent experiments. Data are presented as mean ± SD and analyzed using two-tailed unpaired Student’s *t*-test.

Next, we examined the effect of Zotatifin on cell proliferation by measuring the level of BrdU incorporation in tumors. Zotatifin treatment suppressed DNA synthesis in 2153L, 2225L-LM2, and 2208L tumors in vivo (**Fig. 1E; Supplementary Fig. S1E and F**). To further study the inhibition of cell proliferation, we treated 2153L primary tumor cells in vitro and found that Zotatifin suppressed the G1/S cell cycle transition (**Fig. 1F**). These data indicate that Zotatifin can inhibit proliferation in a tumor cell-autonomous manner.

### Zotatifin suppresses the infiltration of neutrophils and M2-like macrophages and sensitizes tumors to immune checkpoint blockade

In addition to tumor cell-autonomous effects, we interrogated the tumor immune microenvironment in these syngeneic GEM models. To determine whether Zotatifin affects tumor-infiltrating immune cells, we carried out mass cytometry for 2153L tumors that were treated with vehicle or Zotatifin for 7 days in vivo. Zotatifin inhibited the infiltration of the following populations: immuno-suppressive M2-like macrophages (CD11b+, F4/80+, Arginase+, PD-L1+), Arginase+ monocytes that are commonly considered as monocytic myeloid-derived suppressor cells (MDSCs), and neutrophils (CD11b+, Ly6C^lo^, Ly6G+) which are commonly defined as granulocytic MDSCs (**Fig. 2A and B**). The reduction of neutrophil infiltration by Zotatifin was also confirmed by immunostaining for the neutrophil marker S100A8 in 2153L, 2225L-LM2, and 2208L tumor tissues (**Fig. 2C; Supplementary Fig. S2A and B**). Concordantly, we also observed decreased production of Cxcl5 (**Fig. 2D**), the chemokine that stimulates the chemotaxis of neutrophils possessing angiogenic properties (34). On the other hand, Zotatifin promoted the infiltration of the following populations: proinflammatory M1-like macrophages (CD11b+, F4/80+, Arginase-, iNOS+, MHCII+), conventional dendritic cells (CD11b-, CD11c+, CD103+, MHCII+), effector CD4+ T cells (CD4+, FoxP3-), and γδ T cells (CD3+, TCRβ-, CD4-, CD8-, TCRδ+) (**Fig. 2A and B**). Decreased neutrophil infiltration and reduced expression of the immunosuppressive macrophage marker CD206 upon Zotatifin treatment were consistently observed across multiple mouse models as analyzed by flow cytometry (**Fig. 2E and F**), indicating a general effect of Zotatifin on the tumor immune microenvironment.

**Figure 2.**
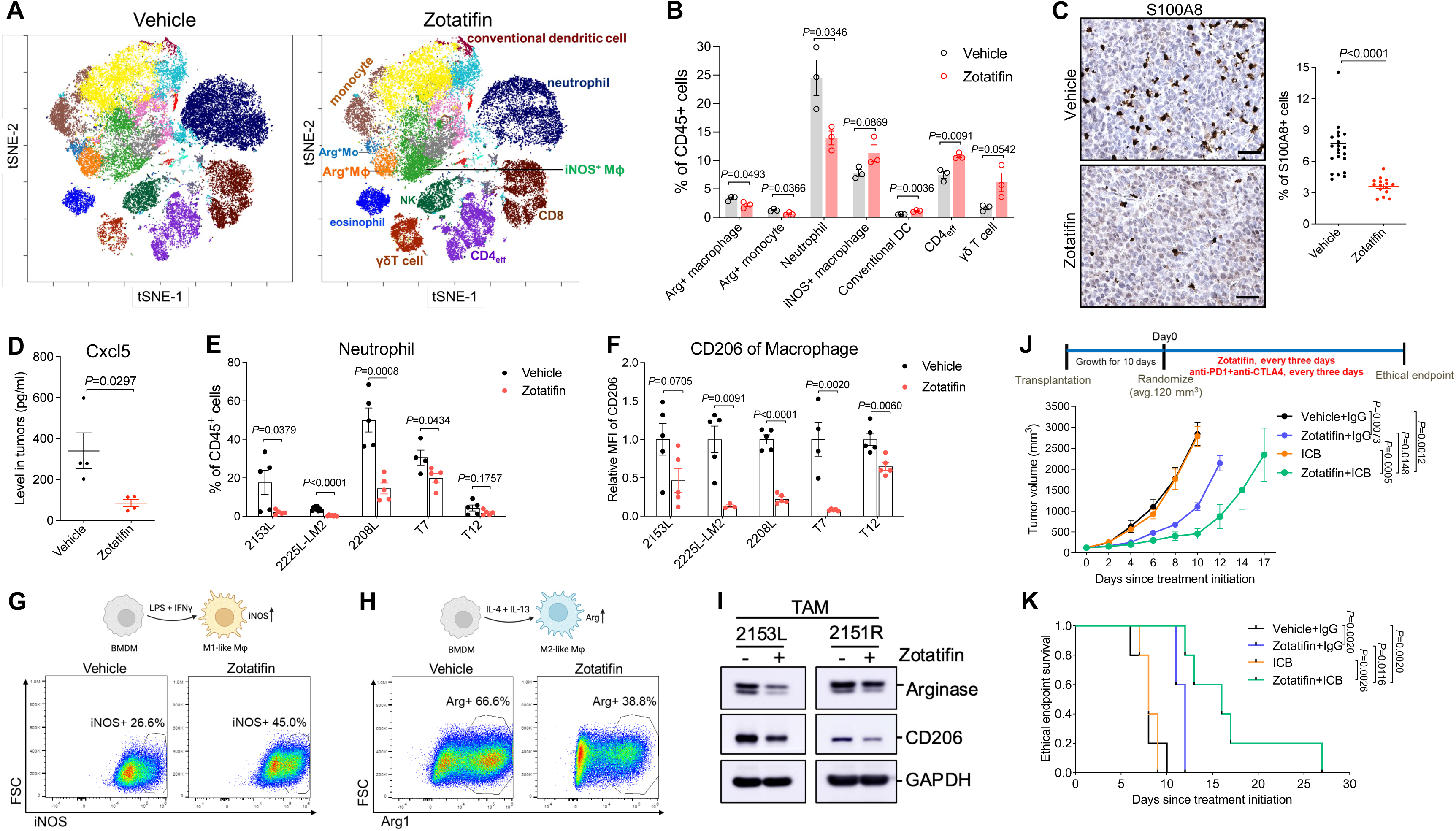
Zotatifin monotherapy alters the tumor immune microenvironment and sensitizes tumors to immune checkpoint blockade. **A,** The t-distributed stochastic neighbor embedding (t-SNE) projection of tumor-infiltrating immune cells of 2153L tumors that were treated with vehicle or Zotatifin for 7 days and analyzed using mass cytometry. Data from three biological replicates of each group were concatenated before t-SNE and FlowSOM clustering analysis. Equal numbers of events are shown for each group and major cell types are marked. **B,** Quantification of major immune cell populations in 2153L tumors that were treated with vehicle or Zotatifin and analyzed using mass cytometry. n=3 biological replicates per group. **C,** Left, representative images of IHC staining of S100A8 in 2153L tumors treated with vehicle or Zotatifin till ethical endpoint. Scale bar, 50 μm. Right, quantification of IHC staining. Three to six representative 20X images were analyzed for each tumor. n=5 biological replicates per group. **D,** Luminex array detection of Cxcl5 levels in lysates of 2153L tumors that were treated with vehicle or Zotatifin for 3 days. n=4 biological replicates per group. **E,** Flow cytometry analysis of tumor-infiltrating neutrophils in ethical endpoint tumors. **F,** Flow cytometry analysis of CD206 median fluorescence intensity (MFI) in tumor-infiltrating macrophages in ethical endpoint tumors. In **E** and **F,** n≥4 biological replicates per group in all models. In **B** to **F**, data are presented as mean ± SEM and analyzed using two-tailed unpaired Student’s *t*-test. **G,** Flow cytometry analysis of iNOS expression in BMDMs that were treated with vehicle or Zotatifin in the presence of LPS and IFNγ. **H,** Flow cytometry analysis of Arginase expression in BMDMs that were treated with vehicle or Zotatifin in the presence of IL-4 and IL-13. **I,** Immunoblotting analysis of TAMs that were separated from treatment naïve 2153L and 2151R tumors and treated with vehicle or 40 nM Zotatifin for 24 hrs in vitro. **J,** Top, treatment scheme of BALB/c mice. Bottom, growth curves of 2153L tumors treated with indicated drugs. n=5 biological replicates per group. Data are presented as mean ± SEM and analyzed using two-way ANOVA with Tukey’s multiple comparison test. **K,** Kaplan-Meier survival curves of 2153L tumor-bearing mice from treatment start time. The log-rank test (two-tailed) was used to test for the significant differences of curves between groups. n=5 biological replicates per group.

We next explored whether Zotatifin can directly repolarize macrophages. When bone marrow-derived macrophages (BMDMs) were cultured in the presence of LPS and IFNγ, the stimuli that induce macrophage polarization towards a M1-like tumor-inhibitory phenotype (35), Zotatifin treatment increased the percentage of iNOS+ cells as compared to the vehicle (**Fig. 2G**). On the other hand, in the presence of IL-4 and IL-13 which are immunosuppressive macrophage polarization stimuli (35), Zotatifin reduced the percentage of Arginase+ BMDMs (**Fig. 2H**). Additionally, we isolated macrophages from untreated 2153L and 2151R tumors and found that Zotatifin treatment suppressed the expression of Arginase and CD206 in these tumor associated macrophages (TAMs) (**Fig. 2I**). These findings suggest that Zotatifin directly promotes the polarization of macrophages towards a tumor-inhibitory phenotype and suppresses their differentiation towards an immunosuppressive phenotype. To confirm the importance of macrophages in the response to Zotatifin monotherapy, we depleted TAMs using an anti-Csf1r antibody (**Supplementary Fig. S2C**) (30). We observed that depleting macrophages partially abolished the response to Zotatifin, suggesting that the repolarization of macrophages contributed to Zotatifin monotherapy response (**Supplementary Fig. S2D**).

High infiltration of immunosuppressive macrophages and neutrophils is associated with a poor response to immune checkpoint blockade (36–40). Given that Zotatifin reduced these immunosuppressive myeloid populations, we hypothesized that it could sensitize tumors to immune checkpoint blockade. We observed that 2153L tumors were completely resistant to CTLA-4 and PD-1 blockade, but the combination with Zotatifin sensitized this immune cold tumor model to checkpoint inhibitors and enhanced survival compared to monotherapy (**Fig. 2J and K**). These data demonstrate that in addition to its tumor-cell autonomous effects, Zotatifin also reprogrammed the tumor immune microenvironment to facilitate the therapeutic benefit of checkpoint inhibition in immune cold TNBC.

### Zotatifin remodels the proteomic landscape and inhibits the translation of *Sox4* and *Fgfr1*

Next, we explored the mechanism underlying the therapeutic effects of Zotatifin monotherapy. Since eIF4A, the major Zotatifin target, primarily affects protein synthesis, we applied quantitative proteomics to unbiasedly investigate changes in steady-state protein levels caused by acute Zotatifin treatment in one of the responsive tumors in vivo. 2153L tumors were treated with either vehicle or Zotatifin for two doses spanning three days before analysis by tandem mass tag mass spectrometry (TMT-MS) analysis (**Fig. 3A**). This treatment design allows for the interrogation of both short-lived and long-lived direct protein targets of Zotatifin, although secondary effects might also be captured. This analysis identified significant alterations in the abundance of 558 of 8531 detected proteins upon Zotatifin treatment that reached the criteria of FDR<0.05, with 333 proteins showing decreased expression and 225 proteins showing increased expression (**Fig. 3B; Supplementary Table S1**). Hallmark gene set enrichment analysis (GSEA) revealed that Zotatifin downregulated the expression of proteins involved in cell cycle progression and stem cell signaling pathways, including E2F targets, G2/M checkpoint, Wnt-β-catenin, and Notch signaling (**Fig. 3B, C, and D; Supplementary Table S2**). On the other hand, proteins involved in interferon α and interferon γ responses were induced in response to Zotatifin treatment (**Fig. 3B, C, and E; Supplementary Table S3**).

**Figure 3.**
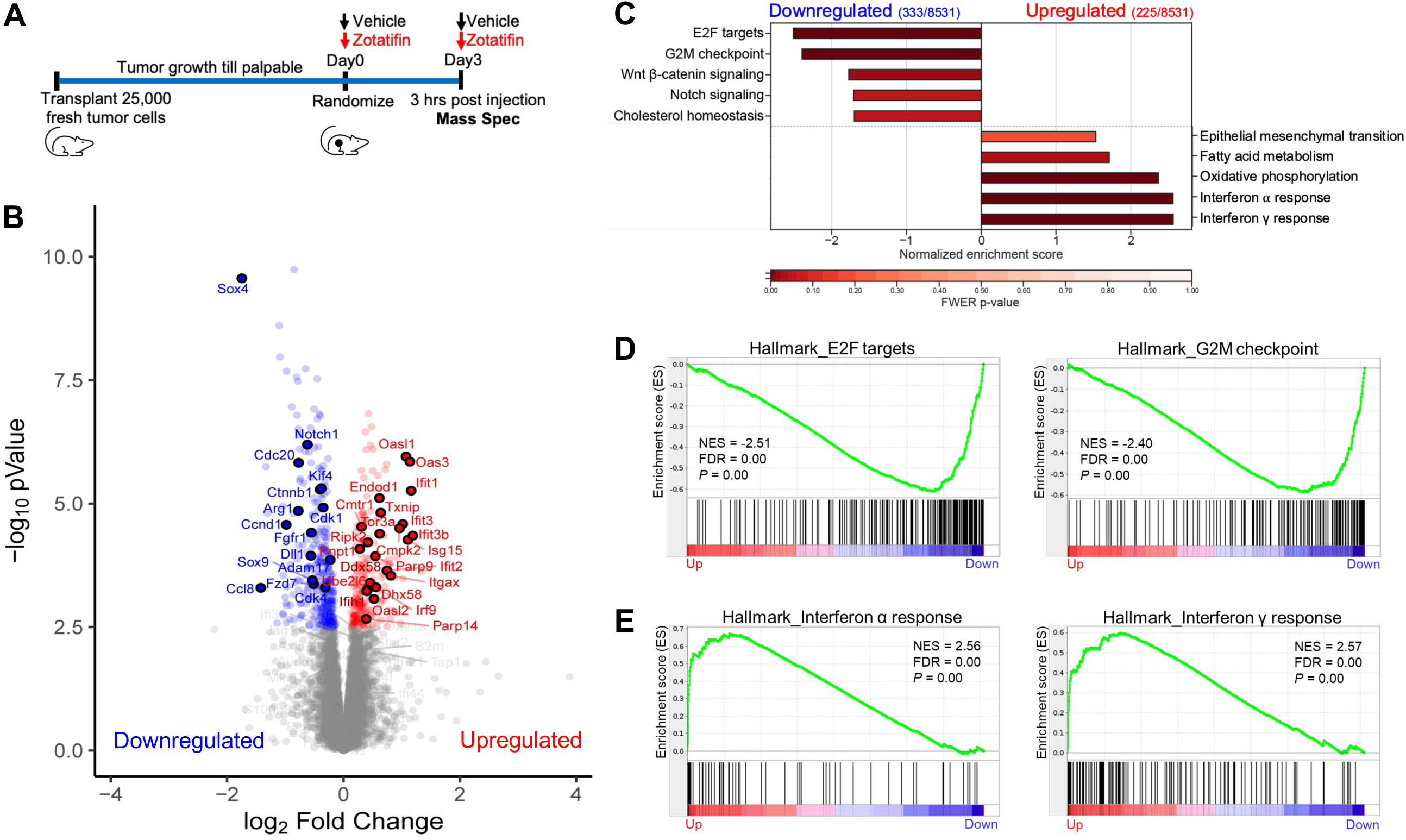
Zotatifin remodels the proteomic landscape of 2153L tumors. **A,** Scheme of sample collection strategy for mass spectrometry. Freshly dissociated 2153L tumor cells were transplanted into the 4^th^ mammary fat pad of BALB/c mice and were allowed to grow until palpable. Mice then were randomized and treated with either vehicle or Zotatifin for two doses spanning 3 days. Tumor tissues were collected 3 hrs after the second injection. **B,** Volcano plot showing relative fold change (log2) in protein abundance versus −log_10_(*P* values) from 2153L tumors treated with Zotatifin compared with vehicle. Proteins that demonstrate a significant change in expression (FDR *q*<0.05) are colored, with decreased expression on the left in blue and increased expression on the right in red. Genes that are critically involved in cell proliferation, stem cell signaling, and interferon response pathways are labeled. n=4 biological replicates per arm. Statistical significance was determined by two-tailed unpaired moderated *t*-test. **C,** GSEA of mass spectrometry results was performed with the MSigDB hallmarks dataset and is summarized as the normalized enrichment score (NES) in vehicle or Zotatifin treated 2153L tumor tissues. Top pathways that have a family-wise error rate (FWER) *P*<0.25 are displayed. FWER *P* values for each pathway are denoted by color. **D,** GSEA enrichment plots for Hallmark E2F targets and G2M checkpoint signatures that are downregulated in Zotatifin treated tumors compared with vehicle. NES, normalized enrichment score. **E,** GSEA enrichment plots for Hallmark interferon α response and interferon γ response signatures that are upregulated in Zotatifin treated tumors compared with vehicle.

We next validated several targets from the proteomic analysis. Sox4 and Fgfr1 were chosen because Sox4 is the most significantly downregulated protein and Fgfr1 is an important receptor tyrosine kinase. First, we performed immunoblotting on a biological replicate of 2153L tumor samples and observed a dramatic reduction of Sox4 and Fgfr1 protein expression in the Zotatifin treatment group (**Fig. 4A**). To determine whether this is a general phenomenon, we performed immunoblotting on the other five GEM models. Strikingly, Zotatifin treatment caused a uniform and potent downregulation of Sox4 and Fgfr1 protein expression in all the GEM models, despite intertumoral variation (**Fig. 4A**). This was also confirmed in 4T1 tumors (**Supplementary Fig. S3A**). Interestingly, Sox4 was not detectable by immunoblotting in the Zotatifin-resistant E0771 although Zotatifin was able to downregulate Fgfr1 expression in this cell line (**Supplementary Fig. S3B**). To investigate at which step this regulation occurs, we quantified *Sox4* and *Fgfr1* RNA expression and found no reduction following Zotatifin treatment in any of the GEM models, except for *Fgfr1* in 2208L (**Fig. 4B and C**), indicating post-transcriptional regulation. Next, we conducted a dose-response analysis of Zotatifin treatment in vitro using primary cells derived from 2153L tumors. We again observed a dose-dependent reduction in Sox4 and Fgfr1 at the protein but not mRNA levels (**Fig. 4D; Supplementary Fig. S3C and D**). A time-course analysis showed that the reduction in both Sox4 and Fgfr1 protein expression was a rapid event that occurred even after 1 hour of exposure to Zotatifin (**Fig. 4E**), consistent with the short half-life of Sox4 and Fgfr1 proteins (**Supplementary Fig. S3E**). The human TNBC cell line BT549 also showed a similar pattern of inhibition of SOX4 and FGFR1 at the protein but not RNA levels upon Zotatifin treatment (**Fig. 4F; Supplementary Fig. S3F and G**). This effect is not due to a nonspecific stress response as Zotatifin did not increase the phosphorylation of eIF2α, an integrated stress response marker, nor did stress-causing chemotherapeutic drugs result in reduction of Sox4 or Fgfr1 (**Supplementary Fig. S3H**). To further explore whether the observed effects of Zotatifin are through targeting eIF4A1, we employed HAP1 cells that express a mutant eIF4A1 (F163L) generated through CRISPR/Cas9-mediated gene editing (41). This mutation can abolish the binding of Zotatifin to eIF4A1 but does not affect the function of eIF4A1 (14). Whereas SOX4 and FGFR1 protein but not RNA expression was downregulated by Zotatifin in wildtype and Cas9 control (F163F) HAP1 cells, these effects were abrogated in eIF4A1-F163L mutant cells (**Fig. 4G; Supplementary Fig. S3I**), demonstrating the specificity of eIF4A1 targeting by Zotatifin. Collectivley these data suggest that the reduction in Sox4 and Fgfr1 protein abundance can serves as robust pharmacodynamic biomarkers for Zotatifin drug activity.

**Figure 4.**
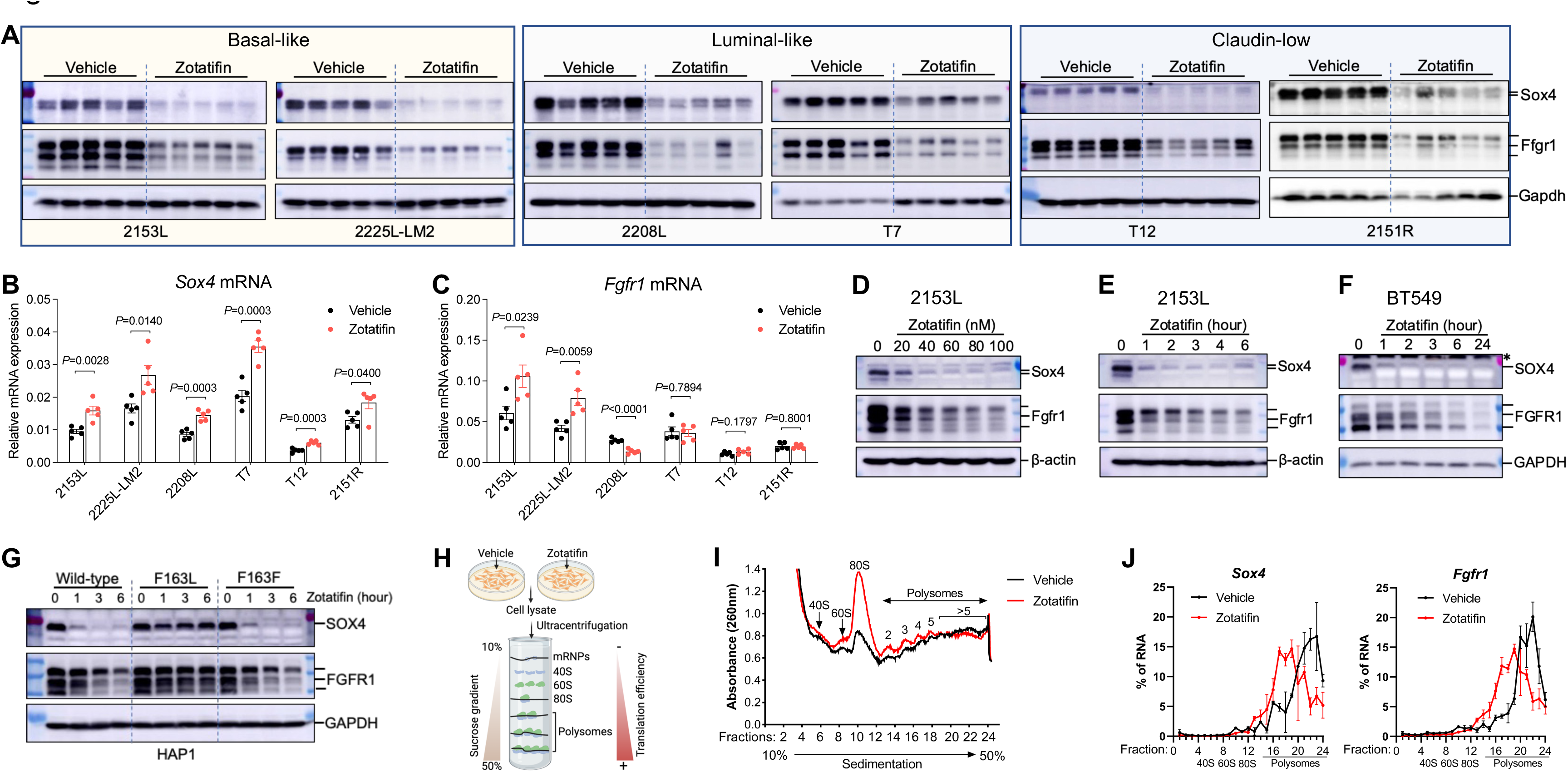
Zotatifin inhibits the translation of *Sox4* and *Fgfr1* mRNAs. **A,** Immunoblotting analysis of tumors that were treated with vehicle or Zotatifin in vivo. n = 5 biological replicates per group. **B** and **C,** QPCR analysis for *Sox4* (**B**) and *Fgfr1* (**C**) mRNA expression in tumors that were treated with vehicle or Zotatifin in vivo. Data are presented as mean ± SEM and analyzed using two-tailed unpaired Student’s *t*-test. n=5 biological replicates per group. **D,** Immunoblotting analysis of 2153L cells that were treated with different concentrations of Zotatifin for 6 hrs in vitro. **E,** Immunoblotting analysis of 2153L cells that were treated with 40 nM Zotatifin for different time periods. **F,** Immunoblotting analysis of BT549 cells that were treated with 40 nM Zotatifin for different time periods. * denotes a non-specific band. In **D**-**F**, data are representative of three independent experiments. **G,** Immunoblotting analysis of HAP1 cells that were treated with 40 nM Zotatifin in vitro. Data are representative of two independent experiments. **H,** Illustration for polysome profiling analysis. **I and J,** Polysome profiling of 2153L cells that were treated with vehicle or 40 nM Zotatifin for 2 hrs. **I,** Representative polysome profiles from three biological replicates. **J**, Distribution of *Sox4* and *Fgfr1* mRNAs across the different fractions. Data are presented as mean ± SEM of three biological replicates.

To further study how Zotatifin regulates Sox4 and Fgfr1 protein expression, we performed polysome profiling of 2153L cells treated with either vehicle or a low concentration of Zotatifin (40nM) for 2 hours in vitro (**Fig. 4H**). This low dose and short time period should mitigate any off-target or secondary effects of drug treatment. We found that Zotatifin treatment dramatically increased the abundance of 80S monosomes, but only had a modest effect on the abundance of polysomes (**Fig. 4I**), suggesting that Zotatifin blocked the translation of a subset of mRNA transcripts. Subsequent qPCR analysis showed that Zotatifin did not affect the translation efficiency of housekeeping genes *Actb* and *Gapdh* (**Supplementary Fig. S3J**); in contrast, it reduced the translation efficiency for both *Sox4* and *Fgfr1* transcripts (**Fig. 4J**). Consistently, cycloheximide (CHX) abolished the Zotatifin-induced reduction of Sox4 and Fgfr1 proteins (**Supplementary Fig. S3K**), demonstrating that Zotatifin inhibits the expression of *Sox4* and *Fgfr1* at the translational level.

### Zotatifin elicits an interferon response through inhibition of *Sox4* translation

The observation of increased expression of proteins involved in interferon α and interferon γ response upon Zotatifin treatment was initially counterintuitive because targeting eIF4A by Zotatifin primarily suppresses mRNA translation. Therefore, we hypothesized that the induction of interferon response related proteins might be a secondary event of eIF4A targeting. This hypothesis was supported by the observation that not only the protein but also the RNA expression of genes such as *Ddx58*, *Ifih1*, and *Tlr3*, which are intracellular pattern recognition receptors (PRRs), and several IFN-stimulated genes (ISGs) were markedly increased upon Zotatifin treatment in 2153L tumors (**Fig. 3B and 5A**). This induction was observed in all six GEM models in vivo (**Fig. 5A**) and in a dose- and time-dependent manner in vitro (**Supplementary Fig. S4A and B**). In contrast, several commonly used chemotherapeutic drugs failed to induce ISG mRNAs (**Supplementary Fig. S4C**). The effect of Zotatifin is specific to targeting eIF4A1 because the induction of interferon response genes was abrogated in eIF4A1-F163L mutant HAP1 cells (**Fig. 5C**). Furthermore, a subset of biopsy samples from heavily pretreated ER+ breast cancer patients treated with Zotatifin in ongoing clinical trials (NCT04092673) showed induction of many of the ISGs compared to pre-treatment (**Fig. 5B; Supplementary Fig. S4D**). Given the heterogenous nature of tumor tissue, these gene expression signals may come from both tumor cells and tumor-infiltrating immune cells. While eIF4A1 is the major Zotatifin target (**Supplementary Fig. S5A and B**), and compared to non-TNBC, TNBCs express higher levels of eIF4A1 at both the RNA and protein levels (**Supplementary Fig. S5C**), unfortunately, no TNBC patients have been included in these trials to date, so a direct comparison with the preclinical data is not possible. Collectively these data indicate that the induction of interferon response RNAs serves as a robust pharmacodynamic biomarker for Zotatifin activity in addition to the reduction of Sox4 and Fgfr1 protein.

**Figure 5.**
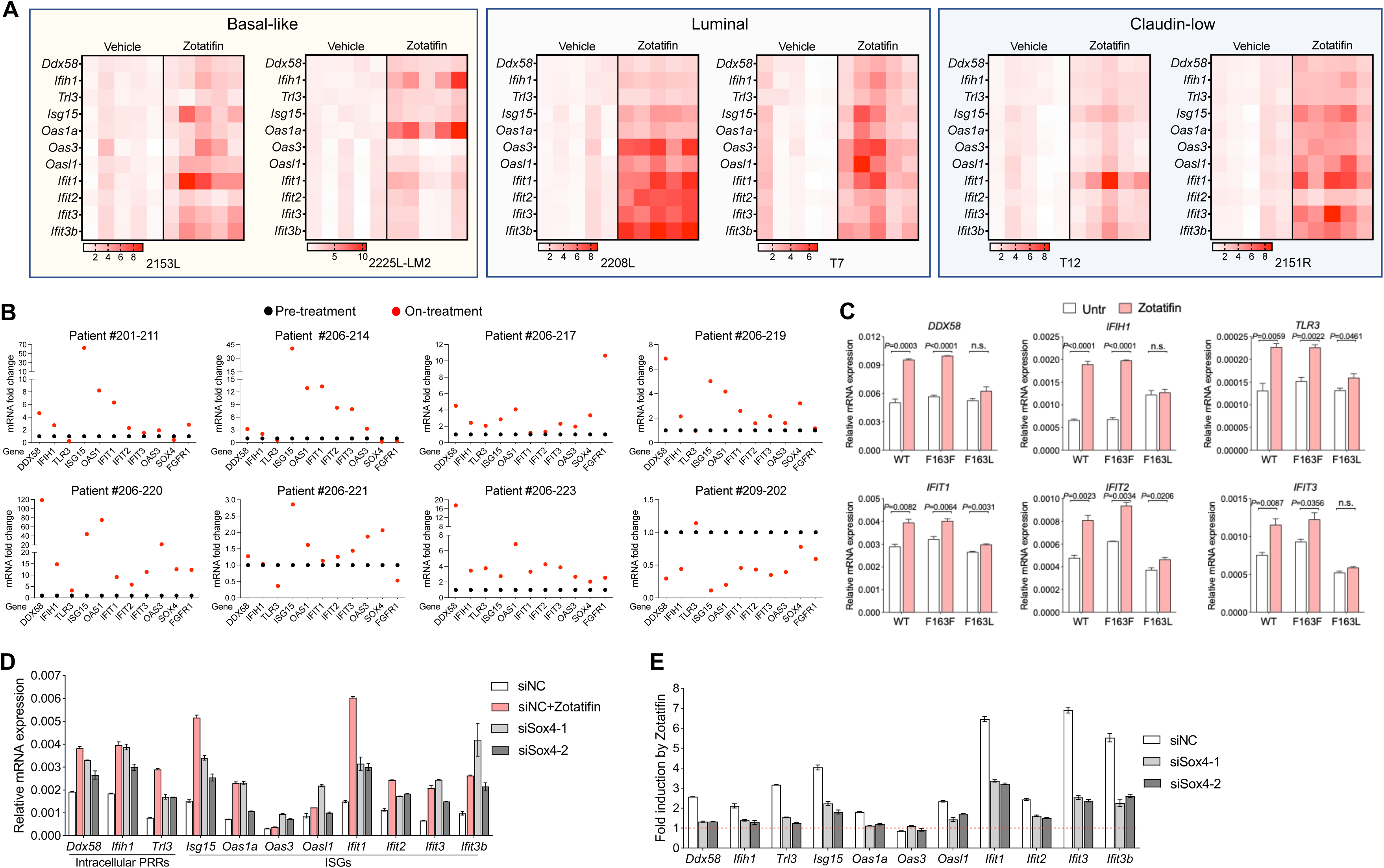
Inhibition of Sox4 translation contributes to Zotatifin-induced interferon response. **A,** QPCR analysis of tumors that were treated with vehicle or Zotatifin in vivo. The mean mRNA levels of the vehicle groups were set as 1 and fold changes were calculated for each gene. n=5 biological replicates per group. **B,** QPCR analysis of 8 paired biopsies from pre-treatment (black) and on Zotatifin treatment (red) ER+ breast cancer patients. The mRNA levels of pre-treatment samples were set as 1 and fold changes were calculated for each paired sample. **C,** QPCR analysis of HAP1 cells that were treated with 40 nM Zotatifin for 6 hrs. **D**ata are representative of two independent experiments and are presented as mean ± SD of technical triplicates. **D,** QPCR analysis of 2153L cells that were transfected with negative control siRNA with or without Zotatifin treatment, or *Sox4* siRNAs without Zotatifin treatment for 48 hrs. **E,** QPCR analysis of Zotatifin-induced gene fold changes in 2153L cells that were transfected with negative control siRNA or *Sox4* siRNAs in the presence of vehicle or Zotatifin. In **D** and **E**, representative data from three biological replicates were shown, and data are presented as mean ± SD of technical duplicates.

Sox4 has been reported to directly suppress the transcription of multiple genes involved in interferon response (42). Accordingly, inhibiting *Sox4* expression using siRNAs (**Supplementary Fig. S5D and E**) increased the levels of PRR and ISG mRNAs in 2153L (**Fig. 5D**) and as a positive control BT549 cells (**Supplementary Fig. S5F**), in which ISGs were previously shown to be regulated by Sox4 (42). Importantly, when *Sox4* was inhibited, the ability of Zotatifin to induce PRRs and ISGs expression was partially impaired (**Fig. 5E; Supplementary Fig. S5G**). Consistently, Zotatifin failed to induce PRR and ISG mRNAs in the Sox4-lacking E0771 cells (**Supplementary Fig. S5H**). These data suggest that Zotatifin induces an interferon response at least in part by inhibiting the translation of *Sox4*.

### Zotatifin synergizes with carboplatin in suppressing tumor growth

Although Zotatifin as a monotherapy was effective in suppressing tumor growth, it did not lead to a durable response. In the clinic, novel targeted therapies will first be tested in combination with standard-of-care therapies. In light of this, we examined whether combining Zotatifin with carboplatin, a routinely used chemotherapy for TNBC (43), would be a more effective treatment. For this purpose, we orthotopically transplanted 2153L tumors into the mammary fat pad of BALB/c mice and initiated either monotherapy or combination treatment when tumors reached 120 mm^3^ (**Supplementary Fig. S6A**). The 2153L tumors were minimally sensitive to carboplatin even at the clinically relevant dose (50 mg/kg); however, the addition of Zotatifin with carboplatin dramatically inhibited tumor growth in four independent experiments (**Supplementary Fig. S6A**) and substantially prolonged survival (**Fig. 6A**). Remarkably, three mice exhibited complete tumor regression following combination therapy and remained tumor-free for months after the treatment stopped.

**Figure 6.**
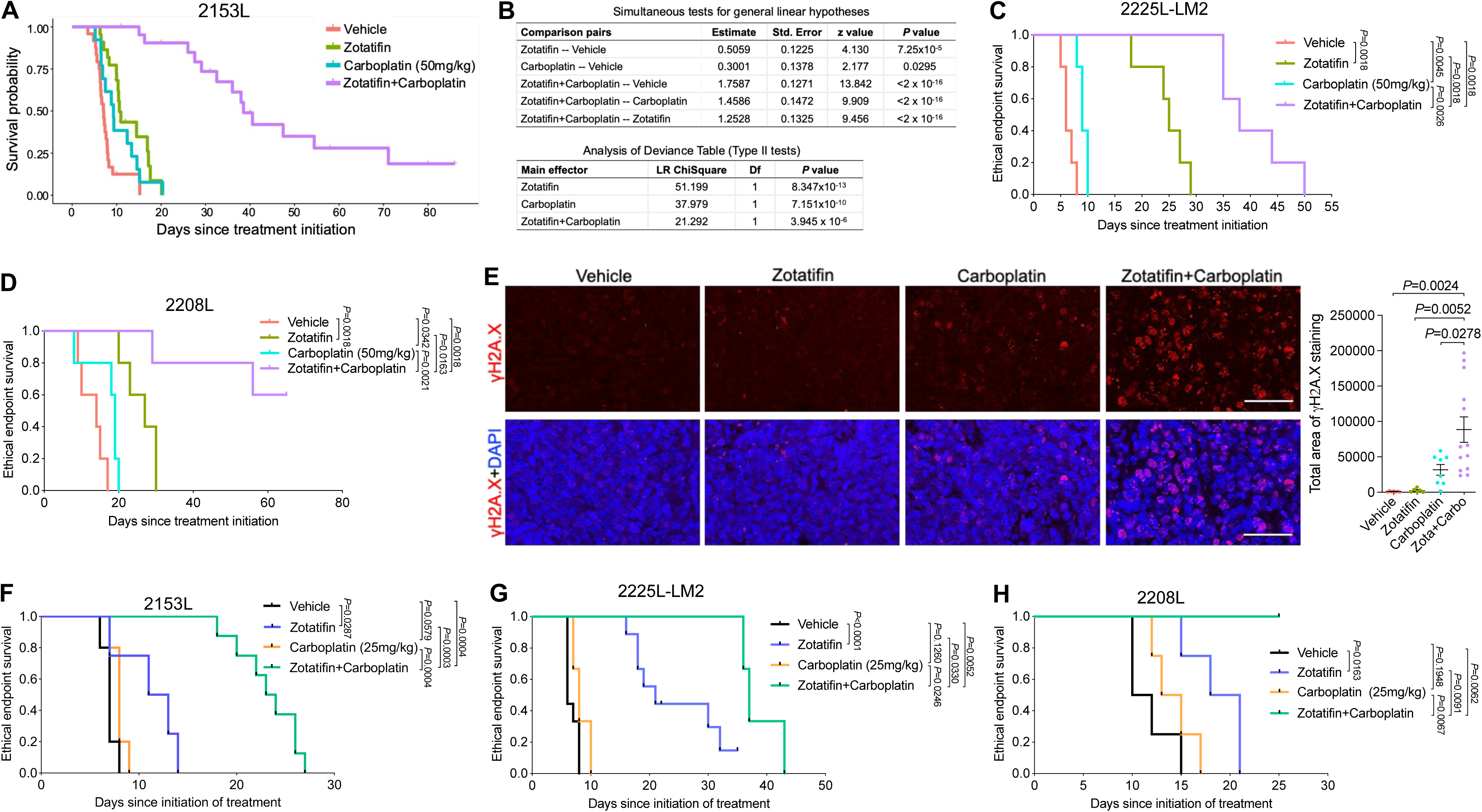
Zotatifin synergizes with carboplatin to suppress tumor progression. **A,** Kaplan-Meier survival curves of 2153L tumor-bearing mice from treatment start time. **B,** Survival regression analysis of 2153L tumor-bearing mice. Survival data are fitted using a parametric survival regression model with a log-normal distribution. Top table reports all possible pairwise comparisons using linear contrasts that are adjusted for multiple comparisons using the Holms method. Bottom table tests for overall main effects and interactions. In **A**-**B**, data from 3 to 5 independent experimental batches are integrated. n=24 for Vehicle, n=22 for Zotatifin, n=13 for Carboplatin, and n=24 for Zotatifin+Carboplatin. **C** and **D,** Kaplan-Meier survival curves of 2225L-LM2 (**C**) or 2208L (**D**) tumor-bearing mice from treatment start time. Mice were randomized and treatment was initiated when 2225L-LM2 tumors reach ∼130 mm^3^ volume or 2208L tumors reach ∼210 mm^3^ volume. n=5 biological replicates per group. **E,** Left, representative images of IF staining of γH2A.X (in red) in 2153L tumors that were treated with indicated drugs for 3 days. Scale bar, 50 μm. Right, quantification of IF staining. At least 2 representative views were analyzed for each tumor and at least 3 tumors for each treatment group were analyzed. Data are presented as mean ± SEM and analyzed using two-tailed unpaired Student’s *t*-test. **F,** Kaplan-Meier survival curves of 2153L tumor-bearing mice treated with indicated drugs. n≥4 biological replicates per group. **G,** Kaplan-Meier survival curves of 2225L-LM2 tumor-bearing mice treated with indicated drugs. n≥3 biological replicates per group. **H,** Kaplan-Meier survival curves of 2208L tumor-bearing mice treated with indicated drugs. n=4 biological replicates per group. In **C**, **D**, and **F-H**, the log-rank test (two-tailed) was used to test for the significant differences of curves between groups.

To assess whether there was a statistical interaction between these two drugs rather than an additive effect of monotherapies, we performed parametric survival regression analysis using the accelerated failure time model (44). This analysis demonstrated that both Zotatifin and carboplatin had a positive effect on mouse survival. However, the combination therapy group had much longer survival than expected based on the additive effect of monotherapies (**Fig. 6B**), indicating a strong interaction between Zotatifin and carboplatin. It is worth noting that the combination therapy was well tolerated and did not cause bodyweight loss (**Supplementary Fig. S6B**). The combination therapy produced a much greater survival benefit than monotherapy in the 2225L-LM2 and 2208L tumor models as well (**Fig. 6C and D**). To investigate the mechanisms responsible for this striking effect, we analyzed 2153L tumors three days after drug treatment. At this early treatment stage, Zotatifin monotherapy had a minimal effect on cell proliferation and apoptosis, while carboplatin monotherapy minimally inhibited cell proliferation but promoted cell apoptosis (**Supplementary Fig. S6C and D**). However, the combination therapy not only notably inhibited cell proliferation but also dramatically induced DNA damage as indicated by the formation of γH2A.X foci (**Fig. 6E)** and enhanced apoptosis (**Supplementary Fig. S6C and D**). These data suggest a strong synergy between Zotatifin and carboplatin.

An important clinical issue is the toxicity of chemotherapies such as neutropenia. If tumors can be sensitized to lower doses of chemotherapy, side effects are likely to be greatly reduced. Therefore, we tested whether Zotatifin could confer a therapeutic benefit to carboplatin at half of the clinically relevant dose (25 mg/kg) in three GEM models. In 2153L, Zotatifin in combination with half-dose carboplatin markedly suppressed tumor growth and prolonged survival compared to monotherapies (**Fig. 6F; Supplementary Fig. S6E**). In 2225L-LM2 and 2208L models, the half-dose of carboplatin had a minimal effect on survival, whereas Zotatifin monotherapy effectively prolonged survival and strikingly in combination with carboplatin led to an overall improved survival benefit (**Fig. 6G and H**). We also tested docetaxel, another routinely used chemotherapy for treatment of TNBC, at half the clinically relevant dose (10 mg/kg) in 2153L. While 2153L was completely resistant to low dose docetaxel, the combination with Zotatifin significantly inhibited tumor growth and prolonged mice survival (**Supplementary Fig. S6F and G**). These data suggest that Zotatifin may effectively sensitize insensitive tumors to low dose chemotherapies.

### Zotatifin synergizes with carboplatin to induce a heightened interferon response and T cell-dependent durable tumor suppression

To investigate the mechanisms underlying the synergistic effect of Zotatifin and carboplatin, we conducted proteomic analysis of 2153L tumors treated with either monotherapy or combination therapy for three days in vivo (**Fig. 7A**). Interestingly, although Zotatifin alone induced a robust interferon response (**Fig. 3C and E**), the combination with carboplatin further markedly increased the interferon response as compared to either monotherapy as revealed in GSEA (**Fig. 7B and C; Supplementary Table S4 and S5**). Combination therapy increased both the number and the level of expression of induced interferon pathway proteins (**Fig. 7D**). Interestingly, this synergy was not observed in vitro as combination treatment did not lead to a greater induction of interferon response genes compared to Zotatifin alone (**Supplementary Fig. S7A**), suggesting that the tumor immune microenvironment contributed to the response. Therefore, we performed mass cytometry of dissociated 2153L tumors and included in the panel Bst2, an interferon stimulated transmembrane protein (45). We observed increased Bst2 expression in almost all the major immune cell populations upon Zotatifin monotherapy and importantly to a greater degree in combination therapy tumors (**Fig. 7E and F; Supplementary Fig. S7B**). These data suggested that Zotatifin and combination therapy not only induced interferon response in tumor cells but also in tumor infiltrating immune cells.

**Figure 7.**
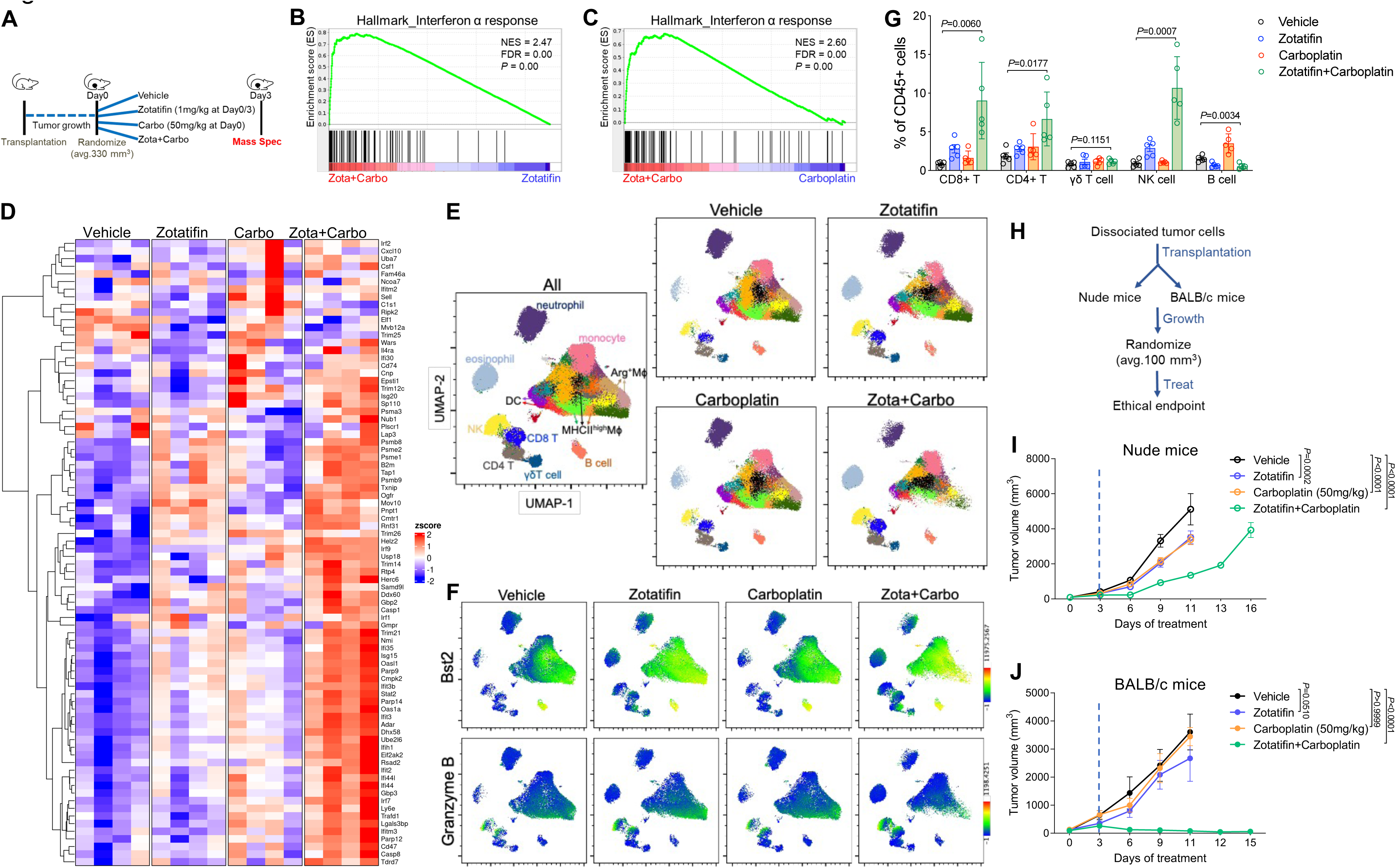
Zotatifin synergizes with carboplatin to induce an interferon response and promote T cell-dependent tumor inhibition. **A,** Scheme of sample collection strategy for mass spectrometry. Freshly dissociated 2153L tumor cells were transplanted into the 4^th^ mammary fat pad of BALB/c mice and were allowed to grow until ∼330 mm^3^ volume. Mice then were randomized and treated with indicated therapies for 3 days. Tumor tissues were collected 3 hrs after the second injection of Zotatifin. n = 4 biological replicates per group. **B,** GSEA enrichment plot for Hallmark interferon α response signature that is upregulated in combination therapy treated tumors compared with Zotatifin monotherapy. **C,** GSEA enrichment plot for Hallmark interferon α response signature that is upregulated in combination therapy treated tumors compared with carboplatin monotherapy. **D,** Heatmap for Hallmark interferon α response signature proteins from tumor tissues treated with indicated therapies. **E-G,** Mass cytometry analysis of tumor-infiltrating immune cells of 2153L tumors that were treated with indicated therapies for 7 days. **E,** The uniform manifold approximation and projection (UMAP) plot overlaid with color-coded clusters. Data from five biological replicates of each group were concatenated before UMAP and FlowSOM clustering analysis. Equal numbers of events are shown for each group and major cell types are marked. **F,** UMAP plot overlaid with the expression of Bst2 or Granzyme B. **G,** Quantification of major lymphoid populations from 2153L tumors in mass cytometry analysis. n=5 biological replicates per group. **H,** Outline of treatment design: freshly dissociated 2153L tumor cells were transplanted into nude mice or BALB/c mice in parallel. Treatment was initiated when tumors reached 100 mm^3^. **I,** Growth curves of 2153L tumors in nude mice treated with indicated drugs. n=5 biological replicates per group. **J,** Growth curves of 2153L tumors in BALB/c mice treated with indicated drugs. n=3 biological replicates for monotherapy groups and n=10 biological replicates for the combination treatment group. In **I** and **J**, data are presented as mean ± SEM and analyzed using two-way ANOVA with Bonferroni’s multiple comparison test.

Combination therapy elicited dramatic changes in both the myeloid and lymphoid compartments of the tumor microenvironment, including decreased infiltration of neutrophils and Arginase+ macrophages and increased infiltration of eosinophils, NK cells, CD8+ T cells, and CD4+ T cells (**Fig. 7E and G; Supplementary Fig. S7B**). In addition, many of the NK cells and CD8+ T cells exhibited Granzyme B expression (**Fig. 7F**). The increased infiltration of CD4+ and CD8+ T cells was also confirmed with immunostaining (**Supplementary Fig. S8A and B**). To investigate whether T cell immunity played a role in the sustained tumor inhibition by combination therapy, the same number of freshly dissociated 2153L tumor cells were transplanted in parallel into immunocompetent BALB/c mice and T cell-deficient athymic nude mice. Treatment was initiated when tumors reached 100 mm^3^ (**Fig. 7H**). In nude mice, Zotatifin and carboplatin monotherapies slowed tumor growth, and combination therapy initially decreased tumor growth for 6 days but was unable to prevent tumor growth beyond this point, and tumors reached the ethical endpoint by day 16 (**Fig. 7I**). In contrast, although monotherapies showed only mild effects in BALB/c mice, the combination therapy elicited marked tumor regression after day 3, which continued at least until day 15 (**Fig. 7J**). These data suggest that while T cells may not affect the efficacy of monotherapy, they contribute to the durable response elicited by combination therapy. Taken together, these findings suggest that inhibition of eIF4A by Zotatifin reprograms the translatome, shifts the tumor immune landscape, and ultimately enhances the response to immune checkpoint blockade or chemotherapy (**Fig. 8**).

**Figure 8.**
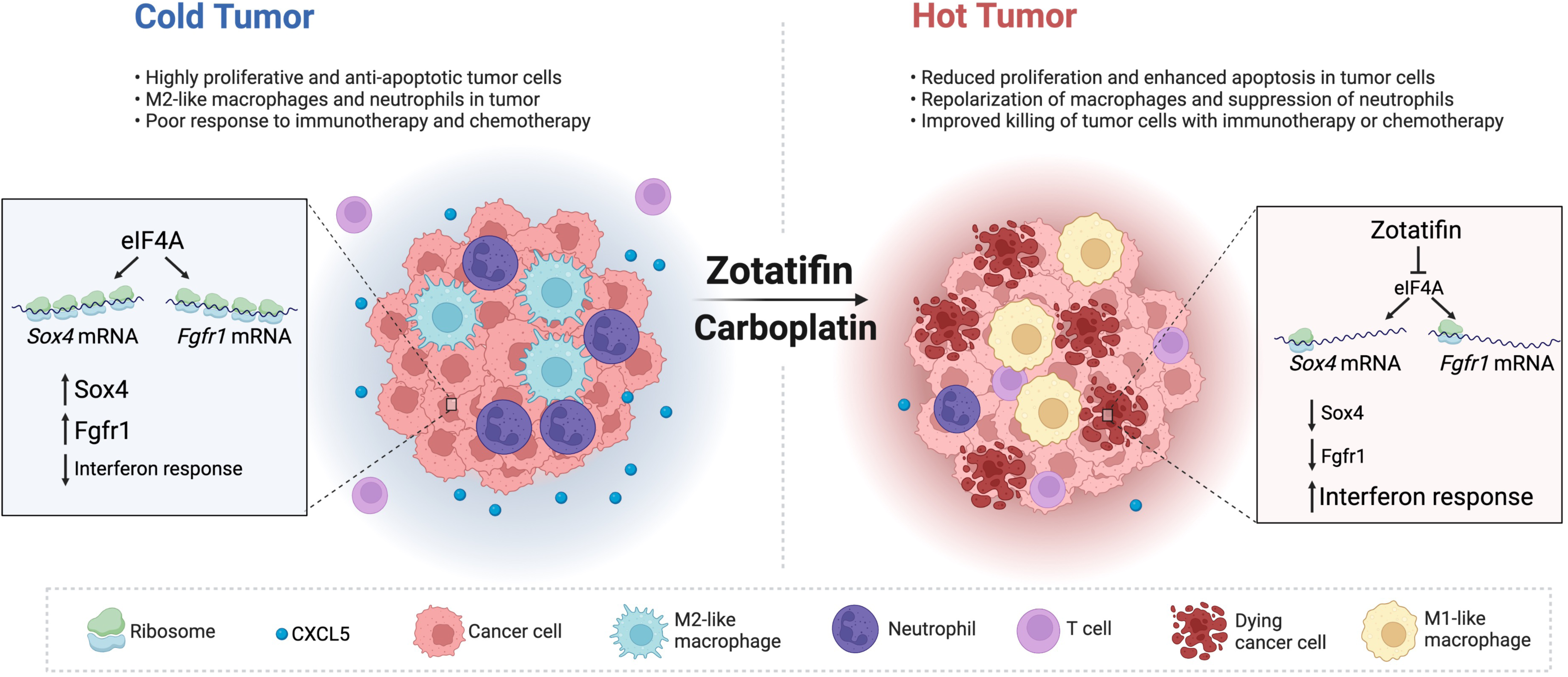
Working model of Zotatifin. Inhibition of eIF4A by Zotatifin suppresses the translation of *Sox4* and *Fgfr1*, induces interferon response, shifts the tumor immune landscape, and ultimately enhances the response to immune checkpoint blockade or chemotherapy.

## Discussion

The RNA helicase eIF4A is a key node where oncogenic signaling pathways converge to impact cancer progression. The present studies demonstrate that targeting eIF4A by Zotatifin can robustly inhibit the translation of *Sox4* and *Fgfr1* in TNBC GEM models, resulting in not only the inhibition of cell proliferation, but also the induction of interferon-related pathways and remodeling of the tumor immune microenvironment. The synergism with carboplatin is exciting as chemotherapies that encompass taxanes, anthracyclines, and platinums remain the standard of care for TNBC patients. Importantly, the inhibition of *SOX4* and *FGFR1* translation by Zotatifin was observed in both mouse tumors and human breast cancer cell lines, indicating the conserved regulation of these genes by eIF4A. The upregulation of a number of ISGs in on-treatment breast cancer patient biopsy samples further suggests that these findings may be translatable from mouse preclinical models to patients.

Mammals possess two highly homologous eIF4A paralogs, eIF4A1 and eIF4A2, that share 90% identity of amino acid sequence and both are targets of Rocaglates. Although functionally similar, they are not redundant. In general, eIF4A1 is much more abundant than eIF4A2 (8), which was also observed in the GEM models used in this study and in human breast cancers from the CPTAC consortium (**Supplementary Fig. S5A and C**) (46). Interestingly, eIF4A1, but not eIF4A2, is a common essential gene in most cancer cell lines in CRISPR knockout screens (**Supplementary Fig. S5B**) (47). Besides eIF4A1 and eIF4A2, a recent study identified DDX3X as another target of RocA (48). Similar to eIF4A1/2, DDX3X is a DEAD-box RNA helicase that has been shown to assist translation initiation in yeast, although its function in translation initiation in mammals is more elusive (49). It is important to note that Rocaglate treatment may not phenocopy the loss of eIF4A activity (13, 50), probably because of its unique mechanism of action. In the case of all the three targets, RocA can increase the RNA affinity of target proteins and clamp the targets onto polypurine RNA in an ATP-independent manner, which inhibits protein synthesis from polypurine-driven reporter mRNAs (48). Importantly, supplementation of recombinant target proteins into an in vitro translation reaction further enhanced RocA-mediated translational repression of polypurine reporter mRNA, indicating a dominant-negative mechanism of action of RocA (15, 48). In contrast, translation inhibition by Hippuristanol, which decreases the affinity between eIF4A and RNA and thus mimics the loss of eIF4A function, was relieved by supplementation with recombinant eIF4A (18, 19). Because RocA converts its targets into dominant-negative repressors, the abundance of its target proteins should predict RocA sensitivity, which has been shown in a few cancer cell lines (48), although this remains to be validated using a larger sample size. Furthermore, it is likely that other genetic and epigenetic differences will affect the treatment response. Clinically, breast cancer is divided into different subtypes based on the expression of different steroid receptors and HER2 as well as the Pam 50 gene signature. Upon interrogating Zotatifin target abundance in different breast cancer subtypes, we found that the protein level of EIF4A1 is higher in TNBC as compared to non-TNBC breast cancer (**Supplementary Fig. S5C**). This suggests that TNBCs may be more sensitive to Zotatifin than receptor-positive breast cancers. The current ongoing clinical trials for Zotatifin in breast cancer (NCT04092673) have been primarily in the context of endocrine resistance in heavily pretreated ER+ patients. These results provide the rationale for testing Zotatifin especially in combination with chemotherapy or immunotherapy in future clinical trials in TNBC patients.

Structural analysis revealed that Rocaglates target a bimolecular cavity formed by eIF4A and polypurine RNA and the Phe163 (F163) on eIF4A1 is crucial for this interaction (14). The specificity of Rocaglates to eIF4A has been extensively addressed by multiple studies. A Rocaglate-resistant eIF4A1 mutant (F163L) has been characterized, and introduction of this allele into cells using CRISPR/Cas9-mediated gene editing confers resistance to Rocaglate cytotoxicity (14, 41, 51, 52). In this study, the near complete abrogation of the effects of Zotatifin on SOX4/ FGFR1/ interferon response genes in eIF4A1-F163L HAP1 cells demonstrated the specificity of Zotatifin to eIF4A1 (**Fig. 4G and 5C**).

The mechanism of action of Rocaglates imposes sequence selectivity onto the general translation factor eIF4A and mRNAs that contain polypurine stretches are more prone to Rocaglates-induced repression of mRNA translation. This selectivity is important because it can potentially lead to lower toxicity and better tolerance in animals and humans. Our observation that only a subset of proteins showed altered translation efficiency in response to Zotatifin in polysome profiling (**Fig. 4I, J and Supplementary Fig. S3J**) further attests to the drug selectivity. Moreover, as normal mouse mammary glands express much lower levels of eIF4A1 than GEM models (**Supplementary Fig. S5A**), this further provided a window of selectivity that contributed to the tolerability of Zotatifin in mice. In the ongoing clinical trial in ER+ breast cancer patients, Zotatifin in combination with either Fuvelstrant or Fulvestrant and Abemaciclib were well tolerated with most adverse effects being grade 1/2 (53). This favorable safety profile is important for its continuous clinical development in potentially other types of cancer. The effect of Rocaglates on tumor cells has been characterized in several studies (10, 12, 13, 20, 21), but their effects on the tumor immune microenvironment remain largely unexplored. An important outcome of our study is the observation of the pleiotropic effects of Zotatifin, impacting both tumor cells and tumor-infiltrating immune cells. This illustrates the importance of drug testing in preclinical immune-competent hosts. Our syngeneic GEM models provide a novel resource of immune-competent animals for these types of therapeutic studies. Comparative oncogenomics and gene profiling have “credentialed” these models and demonstrated that the resulting mammary tumors are representative of the corresponding human breast cancer subtypes (25–27). In addition, we have demonstrated the utility of these preclinical models not only to study single targeted agents, but also to investigate combination treatments with chemotherapy and immunotherapy. A recent study also examined the direct effects of a few Rocaglates including Zotatifin on primary human immune cells and observed various effects (54). Interestingly, Rocaglates have been shown to increase macrophage resistance to *Mycobacterium tuberculosis* infection by directing macrophage polarization toward the M1-like phenotype (55). This is consistent with the results demonstrating the effects of Zotatifin on macrophages both in vitro (**Fig. 2G-I**) and in vivo (**Fig. 2F**). Regarding the mechanism of macrophage polarization, we observed increased Bst2 expression in TAMs upon Zotatifin treatment by mass cytometry (**Fig. 7E and F**). As Bst2 is a type I IFN biomarker (45), we speculated that an induced type I IFN response promoted the polarization of macrophages towards a M1-like phenotype, as reported in multiple studies (56–59). However, this hypothesis awaits further functional studies using Ifnar1/2 knockout macrophages. Due to their immunosuppressive effects, tumor-infiltrating macrophages have been a target for the development of many experimental drugs. Dual targeting of both tumor cells and macrophages is an important property of Zotatifin. Recently, SOX4 was reported to be a critical transcription factor driving the dysfunction of CAR T cells and disruption of SOX4 improved CAR T effector function (60). Given the uniform and potent downregulation of SOX4 by Zotatifin treatment in multiple independent tumor models and cell lines, it will be interesting to explore the possibility that Zotatifin might potentiate CAR T therapy.

The dramatic synergism between Zotatifin and carboplatin is exciting and intriguing. Silvestrol has been shown to enhance sensitivity to doxorubicin by inducing apoptosis in mouse lymphoma models driven by PTEN inactivation or eIF4E overexpression (13). Silvestrol derivatives have also been shown to suppress the expression of CDC6 and induce DNA damage and cell death in a fibrotic pancreatic tumor xenograft model (21). Our study demonstrated that Zotatifin synergizes with carboplatin in mounting a heightened type I interferon response. The type I interferon response can drive growth arrest and apoptosis (61). Indeed, increased DNA damage and cell death was observed after three days of treatment with Zotatifin and carboplatin combination therapy (**Fig. 6E; Supplementary Fig. S6D**). It is likely that this effect is tumor cell autonomous, as T cell immunity might not have occurred at this early stage. The similar tumor growth kinetics observed in nude and BALB/c mice before day 6 support this hypothesis (**Fig. 7I and J**). In contrast to the early response, durable tumor suppression by combination therapy was observed only in BALB/c mice, indicating that T cells contribute to the long-term therapeutic effect. Type I interferon signaling bridges the innate and adaptive immune responses and contributes to therapeutic efficacy (62, 63). Thus, it is possible that the robust interferon response induced by combination therapy in these studies promoted T cell immunity. Besides carboplatin, Zotatifin also synergized with docetaxel (**Supplementary Fig. S6F and G**), a non-DNA damaging chemotherapy routinely used in combination with carboplatin for TNBC treatment (43). It remains to be determined whether Zotatifin in combination with docetaxel will trigger disparate immune responses and whether Zotatifin in combination with both carboplatin and docetaxel will provide greater therapeutic benefit.

Our study has some limitations. First, we recognize that the transplanted syngeneic TNBC models used in the study may not fully capture the complex nature of tumorigenesis over time and its influence on the immune system. Currently it is not feasible to get sufficient numbers of mice with size matched tumor cohorts using autochthonous spontaneous tumor model to study the effects of different therapies both singly and in combination. However, we designed our preclinical studies to include models representing the different molecular subtypes of TNBC and started treatment when tumors were approximately 100mm^3^ in size. Second, we recognize that knockdown of *Sox4* does not fully phenocopy the effect of Zotatifin in inducing ISG expression. Considering the complexity of Zotatifin action, other mechanisms in addition to *Sox4* repression may control ISG abundance. Thirdly, although we showed that both tumor cells and immune cells exhibited induced interferon response upon Zotatifin treatment, we recognize that the source of interferon response genes in heterogenous tumor tissues can only be fully deconvoluted with single cell RNA sequencing and spatial transcriptomics. Finally, although we identified a few pharmacodynamic biomarkers for Zotatifin, the predictive biomarkers for Zotatifin sensitivity remain to be identified.

In summary, strategies to augment ICB and chemotherapy are needed for TNBC treatment. The present studies have identified pharmacodynamic biomarkers for Zotatifin and provided additional preclinical insight into the mechanisms of Zotatifin action both as a single agent and in combination therapies. These studies have provided an important foundation for submission of an investigational new drug application to the FDA to facilitate the next stage of clinical trials for TNBC and potentially other cancer types that also require chemotherapy. Hopefully, these results will help inform future clinical trials and positively impact the treatment of cancer patients.

## Materials and Methods

### Tumor transplantation and treatment

Mice were housed in a room with a 14 hrs light/10 hrs dark cycle at a temperature of 68-72°F and a humidity of 30-70%. The *Trp53*-null GEM TNBC models were previously generated by transplantation of *Trp53*-deleted donor mammary epithelium into the cleared mammary fat pad of syngeneic BALB/c hosts, which gave rise to a cohort of heterogeneous and re-transplantable mammary tumors (24). For tumor transplantation, GEM tumors were first freshly dissociated into single cells as previously reported (64). Briefly, tumor tissues were digested in 1 mg/ml Collagenase A (Sigma #11088793001) for 2 hrs at 37°C with 125 rpm rotation followed by short centrifugation to enrich tumor organoids. Tumor organoids were subsequently trypsinized and filtered into single cells. Then 25,000 tumor cells were transplanted into the 4^th^ mammary fat pad of 6-8-week-old female BALB/c mice (Envigo #047) or Athymic Nude mice (Hsd:Athymic Nude-*Foxn1^nu^,* Envigo #069). For cell line models 4T1 and E0771, 200,000 tumor cells were transplanted into the 4^th^ mammary fat pad of 6-8-week-old female BALB/c mice or C57BL/6 mice (Jackson Laboratory #000664), respectively. When tumors reached an average size of 70-150 mm^3^, mice were randomized followed by drug treatment. Tumor volume and body weight were measured every 2 to 3 days. Investigators were not blinded to the group assignment. Tumor volume was calculated as ½ Length x Width^2^. The ethical endpoint was met when the tumor reached a volume of 1500 mm^3^. For BrdU labelling of proliferative cells, mice were injected with 60 mg/kg BrdU (Sigma #B-5002) 2 hrs before sacrifice.

The following drugs and dosages were used in this study: Zotatifin (eFT226, eFFECTOR Therapeutics) was dissolved in 5% dextrose (Sigma #G7528) and dosed at 1 mg/kg every 3 days; carboplatin (Sigma #C2538) was dissolved in PBS and administered at 25 or 50 mg/kg weekly; docetaxel (LC Laboratories #D-1000) was first dissolved in Tween 80 and then diluted 1:4 with 16.25% ethanol and administered at 10 mg/kg weekly; IgG (BioXCell # BE0089 and # BE0086), anti-PD1 (BioXCell # BE0146), and anti-CTLA-4 (BioXCell # BE0164) were diluted in PBS and dosed at 10 mg/kg every 3 days; Anti-Csf1r (Syndax Pharmaceuticals #SNDX-ms6352) was dosed at 40 mg/kg for the initial dose followed by 20 mg/kg for the remaining dose every week. All drugs and vehicles were administered via intraperitoneal injections.

### Primary cell line generation and cell culture

To establish the primary 2153L cell line, single cells isolated from 2153L tumor tissues were cultured in medium containing 50 μg/ml of G418 (Thermo Fisher #10131035) for two weeks to deplete stromal cells, because the genetic engineering event that led to the deletion of *Trp53* alleles in *Trp53*-null GEM models involved a neomycin resistance gene. Established 2153L cells were cultured in DMEM/F-12 medium (Thermo Fisher #11330032) supplemented with 10% fetal bovine serum (FBS, GenDEPOT #F0900-050), 5 µg/ml insulin (Sigma #I-5500), 1 µg/ml hydrocortisone (Sigma #H0888), 10 ng/ml epidermal growth factor (Sigma #SRP3196), and 1X Antibiotic-Antimycotic (Thermo Fisher #15240062). 2153L cells were tested to be free of mycoplasma contaminants using the Universal Mycoplasma Detection Kit (ATCC #30-1012K). BT549 cells were acquired from ATCC and cultured in RPMI-1640 (GenDEPOT #CM058-050) supplemented with 10% FBS and 1X Antibiotic-Antimycotic. 4T1 and E0771 cells were acquired from Dr. Xiang Zhang’s laboratory at BCM (28) and cultured in DMEM (GenDEPOT #CM002-050) supplemented with 10% FBS and 1X Antibiotic-Antimycotic, HAP1 cells were acquired from eFFECTOR therapeutics (41) and cultured in IMDM (Thermo Fisher #12440046) supplemented with 10% FBS and 1X Antibiotic-Antimycotic. The following drugs were used for cell culture treatment: cycloheximide (CHX, Sigma #C4859) at 100 μg/ml, MG132 (MedChemExpress #HY-13259) at 50 μM, docetaxel (LC Laboratories #D-1000), camptothecin (CPT, MedChemExpress #HY-16560), and doxorubicin (LC Laboratories #D-4000).

In siRNA knockdown assays, 50 µM siRNA (Sigma) was reverse transfected into cells for 48 hrs using RNAiMax (Thermo Fisher #13778030). The following siRNAs were used in this study: si-mmu-Sox4-1 (Sigma #SASI_Mm01_00114970), si-mmu-Sox4-2 (Sigma # SASI_Mm01_00114972), si-has-SOX4 (Sigma # SASI_Hs01_00188751) and siRNA Universal Negative Control #1 (Sigma # SIC001).

### Separation of tumor-associated macrophage (TAM)

Untreated 2153L and 2151R tumors were dissociated in 1 mg/ml Collagenase A for 2 hrs at 37°C and TAMs were separated using EasySep™ Mouse F4/80 Positive Selection Kit (STEMCELL technologies, #100-0659) following manufacture’s protocol. TAMs were cultured in BMDM culture medium and treated with either vehicle or 40 nM Zotatifin for 24 hrs before immunoblotting.

### Patient biopsy samples

De-identified frozen core biopsy samples were obtained from 8 ER+ breast cancer patients who were heavily pre-treated and then enrolled in the Zotatifin Phase I/II clinical trial (NCT04092673). Tumor tissues came from either primary (for patient #201-211) or metastatic (for the additional 7 patients) sites. All the patients received 0.07 mg/kg of Zotatifin on a 2 weeks on/1 week off schedule. Patient #206-214 received Zotatifin monotherapy and the additional 7 patients received Zotatifin in combination with fulvestrant. Paired pre-treatment and on-Zotatifin treatment biopsies were acquired from each patient, and on-treatment specimens were collected at C1D9 (approximately 24 hrs after the second dose of Zotatifin).

### Immunohistochemistry (IHC) and immunofluorescence (IF)

Tumor tissue specimens were fixed in 4% PFA overnight and stored in 70% ethanol until paraffinization and embedding. Tissue sections of 5 μm thickness were deparaffinized, rehydrated, and subjected to antigen retrieval in citrate buffer (pH 6.0, Sigma #C9999) for 20 min in a steamer. Slides were incubated with primary antibodies overnight at 4°C and with secondary antibodies for 1 hr at room temperature (RT) before ABC-HRP (Vector Laboratories #PK-7100) treatment and DAB (Vector Laboratories #sk-4105) development for IHC or DAPI staining for IF. The antibodies and dilutions used were as follows: F4/80 (1:1000, Cell Signaling Technology (CST) #70076S), S100A8 (1:5000, R&D Systems #MAB3059), BrdU (1:1000, Abcam #ab6326), cleaved Caspase3 (1:1000, CST #9661), phospho-Histone H2A.X (Ser139) (1:400, CST #9718), CD4 (1:800, Abcam #ab183685), and CD8 (1:800, eBioscience #14-0808-82).

All IHC slides were scanned with Aperio and analyzed using the Aperio ImageScope software (v12.3.3.5048). At least 3 representative 20X views were analyzed for each tumor and at least 3 tumors for each treatment group were analyzed. Specifically, the percentage of BrdU+, cleaved Caspase3+, or CD4+ cells was analyzed using the Nuclear V9 algorithm. The total percentage of positive nuclei was counted (including strongly, moderately, and weakly stained cells) for BrdU+ and cleaved Caspase 3+ cells, and only the strongly stained signals were counted for CD4+ cells. To calculate the percentage of S100A8+ or CD8+ cells, positively stained cells were counted manually to avoid the inclusion of background staining. This number was then divided by the total cell number for each view, which was automatically counted using the Aperio Nuclear V9 algorithm. IF images were taken using Nikon A1-Rs confocal microscope at 40X and analyzed using Python (version 3.8.15). Specifically, the γH2A.X channel mask is obtained by first applying the MaxEntropy thresholding filter followed by applying a binary closure with Gaussian filter (sigma=2). Any object smaller than 9 pixels is removed to obtain the final mask. At least 2 representative views were analyzed for each tumor and at least 3 tumors for each treatment group were analyzed.

### Mass cytometry and flow cytometry

*Trp53*-null tumors treated with vehicle or Zotatifin were dissociated in 1 mg/ml Collagenase A for 2 hrs at 37°C with 125 rpm rotation followed by 3 short centrifugations to enrich for tumor stromal cells from supernatants. Red blood cells were removed using RBC lysis buffer (Biologend #420301). The remaining single cell suspension was analyzed using mass cytometry or flow cytometry.

In mass cytometry, after staining with cisplatin viability dye, cells were surface stained using a cocktail of antibodies conjugated to metal isotopes, fixed, permeabilized using Foxp3 staining buffer set (eBioscience #00552300), and stained with antibodies for intracellular markers. The cells were then stained with Cell-ID Intercalator-Ir (Fluidigm #SKU201192A) overnight at 4°C and analyzed using a Helios CyTOF Mass Cytometer (Fluidigm). The normalized FCS files were first processed using FlowJo to remove beads, dead cells, and doublets. Equal numbers of CD45+ single cells from biological replicates of each treatment group were concatenated and subjected to analysis in Cytobank. Data were dimensionally reduced using viSNE and cell clusters were identified using FlowSOM.

For flow cytometry of tumor-infiltrating immune cells or BMDMs, single cells were stained with LIVE/DEAD Fixable Yellow Dead Cell Stain (Thermo Fisher #L34967), blocked with anti-CD16/32 (Biolegend #101330), and stained with surface and intracellular markers according to manufacturer’s instructions of the Foxp3 staining buffer set. The following antibodies were used for flow cytometry: CD45 (Biolegend #103128), CD11b (Biolegend #101227), CD206 (Biolegend #141732), Ly-6C (Biolegend #128017), Ly-6G (Biolegend #127651), F4/80 (Biolegend #123115), iNOS (Miltenyi Biotec #130116421), and Arginase (Invitrogen #12369782).

For cell cycle analysis, 2153L cells cultured in regular medium in vitro were allowed to grow to 30% confluence and treated with 40 nM Zotatifin for 48 hrs before trypsinization and fixation in 4% PFA. Next, the cells were pelleted, washed with PBS, and resuspended in DAPI staining buffer (Thermo Fisher #R37606) before flow cytometry.

All flow cytometry data were acquired using an Attune NxT Flow Cytometer (Thermo Fisher) and analyzed using FlowJo software (version 10.7.1). The cell cycle distribution was analyzed using the Watson Pragmatic algorithm provided by FlowJo.

### Luminex cytokine analysis

2153L tumor tissues from the same batch with TMT MS were pulverized to powder under liquid nitrogen and lysed in MILLIPLEX MAP Lysis buffer (Millipore #43040). Lysates were cleared by centrifugation at 10,000 g three times at 4°C. The protein concentration of the supernatant was measured using a BCA Protein Assay Kit and adjusted to 1 mg/ml using lysis buffer. The abundance of cytokines was determined using a MILLIPLEX Mouse 32-Plex Cytokine Panel 1 kit (Millipore # MCYTMAG-70K-PX32) according to the manufacturer’s instructions.

### Tandem mass tag mass spectrometry (TMT MS) and data processing

Sample preparation for deep scale proteomic profiling was performed as described previously by the CPTAC consortium (65) with slight variations. Briefly, frozen tumor tissues were crushed to powder and lysed with urea lysis buffer containing 8 M urea (G-Biosciences #BC89), 75 mM NaCl, 50 mM Tris pH 8.0, 1 mM EDTA (Sigma #E7889), 2 μg/ml Aprotinin (Sigma #A6103), 10 μg/ml Leupeptin (Roche #11017101001), 1 mM PMSF (Sigma #93482-50ML-F), 10 mM NaF, phosphatase inhibitor cocktail 2 (Sigma #P5726), and phosphatase inhibitor cocktail 3 (Sigma #P0044). Samples were lysed for 30 min on ice followed by 2 cycles of 5 sec ON/10 sec OFF sonication (Sonics Materials #GE 505). The lysates were cleared by centrifugation, and the protein concentration was measured using a NanoDrop spectrophotometer (DeNovix #DS-11). Total protein (150 μg) from each sample was reduced with 5 mM dithiothreitol (DTT) for 1 hr at 37°C and alkylated with 10 mM iodoacetamide (Sigma #I1149) for 45 min in the dark at RT. The samples were then diluted with 50 mM Tris pH 8.0, and subjected to Lysyl EndopeptidaseR (Wako #129-02541) digestion for 2 hrs and trypsin (Thermo Fisher #90057) digestion overnight at RT. The digest was acidified with 1% formic acid (Fisher #A117-50) and desalted using Sep-Pak Vac 1cc C18 cartridges (Waters #WAT054955). Elutes were dried with SpeedVac (Thermo Fisher #SC210A) and dissolved in 50 mM HEPES pH 8.5 buffer (Alfa Aesar #J63218).

For TMT labeling, 120 μg digested peptide from each sample as well as RefMix, which is a mixture of equal amounts of peptide from each sample, were labeled with the TMT10plex Label Reagent Set (Thermo Fisher #90110) for 1 hr at RT. After confirming the labeling efficiency for each channel using quality control MS runs, the reaction was quenched by adding 5% hydroxylamine (Thermo Fisher #90115) for 15 min at RT. All samples were then combined and freeze dried using SpeedVac. Each plex was reconstituted with 1 ml of 3% ACN/0.1% formic acid and desalted using Sep-Pak Vac 3cc tC18 cartridges (Waters #WAT054925). The elute was dried using SpeedVac.

TMT-labeled peptides were fractioned offline using an Agilent 300Extend-C18 column (4.6 mm X 250 mm, 5 µm) coupled to an Agilent 1260 Infinity II system at 1 ml/min for 96 min. The 96 fractions were concatenated into 24 peptide pools and a flow-through pool and acidified with 0.1% formic acid. The peptides were separated on an online nanoflow Easy-nLC-1200 system (Thermo Fisher) coupled to an Orbitrap Fusion Lumos ETD mass spectrometer (Thermo Fisher). Proteome fractions (1 µg each) were loaded onto pre-column (2 cm x 100 µm I.D.) and separated on in-line 5 cm x 150 µm I.D. column (Reprosil-Pur Basic C18aq, Dr. Maisch) equilibrated with 0.1% formic acid. Peptide separation was performed at a flow rate of 750 nl/min over a 90 min gradient time with different concentrations of solvent B (4-32% for 85 min, followed by 5 min wash at 90% B). The peptides were ionized at a positive spray voltage of 2.4 kV and the ion transfer tube temperature was set at 300^°^C. The mass spectrometer was operated in data-dependent mode with 2 sec cycle time. MS1 was acquired in Orbitrap (60000 resolution, scan range 350-1800 m/z, AGC 5e5, 50 ms injection time), followed by MS2 in Orbitrap (50000 resolution, AGC 1e5, 105 ms injection time, HCD 38%). Dynamic exclusion was set to 20 sec and the isolation width was set to 0.7 m/z.

To process the proteomics data, raw files were converted to mzML using MSConvert (66). MASIC (67) was used to calculate precursor ion intensities (derived from the area under each elution curve) and to extract reporter ion intensities using default high resolution MS parameters. The Butterworth smoothing method was used with a sampling frequency of 0.25 and an SIC tolerance of 10 ppm. Reporter ion tolerance was set to 0.003 Da with reporter ion abundance correction enabled. Raw spectra were searched with MSFragger (v3.2) using both mass calibration and parameter optimization (68, 69). Peptide validation was performed using a semi-supervised learning procedure in Percolator (70) as implemented in MokaPot (71). The peptides were grouped and quantified into gene product groups using gpGrouper (72). Only the gene products identified in both TMT multiplexes were retained for downstream analyses. Samples were first normalized to the internal reference within each TMT multiplex and then normalized by their median peptide abundance before subsequent data analyses.

### Gene set enrichment analysis (GSEA)

GSEA (v3.0) was performed using hallmark gene sets (v7.0) from the Molecular Signature Database (MSigDB) using default settings after mapping mouse genes to their human homologs using the HomoloGene system. Genes without mapping were excluded, and the median value was taken when multiple mouse genes mapped to a single human gene. Pathway enrichment *P* values were calculated using gene set permutation.

### Immunoblotting assay

Tumor tissues were snap frozen upon harvest and homogenized in lysis buffer (Tris-HCl pH 6.8, 62.5 mM; SDS, 2%; protease inhibitor cocktail (Sigma #11873580001); phosphatase inhibitors (Sigma #4906845001)) using zirconium beads (Benchmark Scientific #D1132-30) in a bead homogenizer. Protein concentrations were measured using the BCA Protein Assay Kit (Thermo Fisher #23227). Whole cell extracts were separated by sodium dodecyl sulfate-polyacrylamide gel electrophoresis and transferred to polyvinylidene difluoride membranes (Millipore #IPVH00010). The antibodies and dilutions used were as follows: Arginase-1 (1:1500, CST #93668), CD206 (1:1000, CST #24595T), FGFR1 (1:1500, CST #9740S), Sox4 (1:4000, Diagenode #C15310129), GAPDH (1:4000, CST #2118), and β-actin (1:4000, CST #3700S). GAPDH and β-actin served as loading controls.

### Quantitative real-time PCR (qPCR)

Total RNA was extracted using TRIzol reagent (Thermo Fisher #15596026) following the manufacturer’s instructions. Total RNA (1 µg) was converted to cDNA using the High-Capacity cDNA Reverse Transcription Kit (Thermo Fisher #4368814). The mRNA levels were detected using amfisure qGreen Q-PCR Master Mix (GenDEPOT #Q5602). *18S* was used as the internal reference gene for patient biopsy samples and *Actb* was used as the internal reference for all other samples. The levels of target genes were normalized to those of internal reference gene to calculate the 2^-ΔΔCt^ value. The sequences of all the qPCR primers are listed in **Supplementary Table S6**.

### Polysome profiling analysis

Sucrose gradients (10-50%) were prepared in polysome buffer (20 mM Tris pH 7.4, 150 mM NaCl, 5 mM MgCl2, 1 mM DTT, 100 μg/ml CHX supplemented with 20 U/ml SUPERaseIn RNase Inhibitor (Invitrogen #AM2696) and RNase-free sucrose (Sigma #84097), poured into polypropylene tubes (Beckman Coulter #331374) the evening before use, and stored at 4°C.

2153L cells were seeded in 15 cm plates overnight and reached 80% confluence on the day of harvest. Cells were treated with DMSO (0.0004%) or 40 nM Zotatifin for 2 hrs before incubation in 100 μg/ml CHX for 5 min. Next, cells were washed with PBS and trypsinized to single cells in the presence of 100 μg/ml CHX, followed by washing in ice-cold PBS + CHX and lysing in 500 μl of ice-cold lysis buffer (polysome buffer supplemented with 1% Triton X-100 and 25 U/ml Turbo DNase I (Invitrogen #AM2238)). Cell lysates were triturated five times through an 18-gauge needle and ten times through a 27-gauge needle, followed by incubation on ice for 10 min. Then lysates were cleared by centrifugation for 10 min at 14,000 g at 4°C, and an equal volume of supernatant was carefully layered on top of the sucrose gradients. The gradients were ultracentrifuged at 35,000 rpm for 2 hrs at 4°C using an SW41 Ti rotor (Beckman Coulter). The gradients were then displaced into a UA-6 continuous UV detector (Teledyne ISCO) using a syringe pump (Brandel) containing Fluorinert FC-40 (Sigma #F9755) at a speed of 0.75 ml/min. Absorbance was recorded at an OD of 260 nm using the Logger Lite (version 1.9.4) software. A total of 24 fractions with 500 μl volume were collected for each sample using the Foxy Jr fractionator.

A volume of 250 μl fraction was mixed with 500 pg of luciferase RNA spike-in (Promega #L4561) and lysed in 750 μl Trizol LS (Thermo Fisher #10296028). RNA was precipitated in the presence of GlycoBlue (Invitrogen #AM9515) and dissolved in 20 μl RNase-free water. Then, 6 μl of RNA from each fraction was converted to cDNA using the High-Capacity cDNA Reverse Transcription Kit (Thermo Fisher #4368814). The cDNA was diluted 10 times before qPCR analysis. The qPCR data was processed as previously reported (73).

### Statistical analysis

Unpaired two-tailed Student’s *t*-tests were performed to compare the differences between two groups in most studies where indicated. For proteomics data, differential analysis was performed using the moderated *t*-test, as implemented in limma (74). Two-way ANOVA with Bonferroni or Tukey’s multiple comparison test was used to analyze tumor volume or body weight over time. The log-rank test was used to test for significant differences in the Kaplan-Meier survival curves between groups for Fig. 2K, 6C, 6D, 6F-H and Supplementary Fig. S6G. Mice whose tumors had never reached the target size were censored at their last time point. Censored events were marked as vertical ticks on the curves. All above analyses were performed using GraphPad Prism 9 software. The survival regression analysis shown in Fig. 6A and 6B was performed using R software. For these data, the time to the endpoint tumor size (1500 mm^3^) for each mouse was computed by linear interpolation using the two time points and tumor sizes immediately before and immediately after crossing the boundary for the first time. Survival data were modeled using a parametric survival regression based on a log-normal distribution with main effects for each treatment and their interaction. All possible pairwise comparisons were computed using linear contrasts and adjusted for multiple comparisons using Holm’s method. All *P* values were two-sided.

### Study approval

All mouse studies were conducted in compliance with the Guide for the Care and Use of Laboratory Animals of the National Institutes of Health. All mice were maintained and euthanized according to the guidelines of IACUC at Baylor College of Medicine (Protocol AN-504). De-identified patient biopsy study was reviewed and approved by IRB at Baylor College of Medicine and was considered as a human material study.

### Data and materials availability

All data generated in this study are available within the article and its supplementary information and from the corresponding author upon reasonable request. All raw mass spectrometry data were deposited in the publicly available MassIVE/ProteomeXchange under accession number MSV000089580/PXD034250.

### List of Supplementary Materials

Supplementary Fig.S1 to S8 Supplementary Table S1 to S6

## Acknowledgments

We thank Dr. Kevin Roarty at BCM for providing the 2225L-LM2 tumor model. We thank Dr. Mahnaz Janghorban for her help with immunostaining. We are grateful to Dr. Chad Creighton at BCM for his help with bioinformatic analyses. We thank Drs. Licheng Zhang, Yitian Xu, and Shu-Hsia Chen at HMRI for their help with mass cytometry. We thank Mei Leng and Antrix Jain at the BCM proteomics core, Dr. Fabio Stossi at the BCM integrated microscopy core, and Drs. Xuan Wang and Shixia Huang at the BCM Luminex core for their technical assistance. We thank Dr. Peter Ordentlich from Syndax for providing the anti-Csf1r antibody. We are grateful to patients who participated in Zotatifin clinical trials, and we thank Dr. Funda Meric-Bernstam at MDACC and Dr. Ezra Rosen at MSKCC for providing patient biopsy samples.

## Funding

This project was supported by the NIH (CA16303-46), CPRIT (RP220468), and eFFECTOR therapeutics to J.M.R. The BCM Mass Spectrometry Proteomics Core was supported by the Dan L. Duncan Comprehensive Cancer Center Award (P30CA125123), CPRIT Core Facility Support Awards (RP170005 and RP210227), and Intellectual Developmental Disabilities Research Center Award (P50HD103555). This project also was supported by the Cytometry and Cell Sorting Core at Baylor College of Medicine with funding from the CPRIT Core Facility Support Award (CPRIT-RP180672), the NIH (CA125123 and RR024574), and the assistance of Joel M. Sederstrom. S.G.H. was supported by CCSG P30CA125123 and SPORE P50CA186784. C.M.P. was supported by R01-CA148761 and SPORE P50CA058223. Illustrations were created with BioRender.com.

## Author contributions

N.Z. conceived the study, designed and conducted the experiments, analyzed the data, and wrote the manuscript. E.B.K., X.Y., L.C.R., N.L., Y.G., D.P., S.C., A.S. and C.H. performed experiments. A.B.S. and A.M. designed the mass spectrometry experiments, analyzed the data, and edited the manuscript. K.S. analyzed the IF images. S.S. and B.Z. analyzed the CPTAC data. J.Z. and L.M.S. scanned immunostaining slides using Aperio. S.G.H. performed biostatistical analyses. C.F. performed bioinformatics analyses. C.M.P. performed bioinformatics analyses and edited the manuscript. J.M.R. conceived and supervised the study and edited the manuscript.

## Competing interests

J.M.R. received research support from eFFECTOR therapeutics. C.M.P. is an equity stockholder and consultant of BioClassifier LLC; C.M.P. is also listed as an inventor on patent applications for the Breast PAM50 Subtyping assay.

## Supplementary figure legends

**Supplementary Figure 1.** Zotatifin inhibits tumor growth in a cohort of *Trp53*-null preclinical models. **A,** Individual tumor growth curves of BALB/c mice treated with either vehicle or Zotatifin. **B,** Body weight changes of tumor-bearing BALB/c mice over the treatment course. In **A** and **B**, n=6 biological replicates for 2225L-LM2 and n=5 for all other models in each treatment arm. **C,** Growth curves of 4T1 and E0771 tumors treated with either vehicle or Zotatifin. **D,** Body weight changes of 4T1 and E0771 tumor-bearing mice over the treatment course. In **C** and **D**, n=5 biological replicates for both models. In **A-D**, data are presented as mean ± SEM and analyzed using two-way ANOVA with Bonferroni’s multiple comparison test. **E** and **F,** Left, representative images of IHC staining of BrdU in ethical endpoint 2225L-LM2 (**E**) or 2208L(**F**) tumor tissues. Scale bar, 50 μm. Right, quantification of IHC staining. Five representative 20X images were analyzed for each tumor. Data are presented as mean ± SEM and analyzed using two-tailed unpaired Student’s *t*-test.

**Supplementary Figure 2.** Zotatifin alters the tumor immune microenvironment. **A** and **B,** Left, representative images of IHC staining of S100A8 in 2225L-LM2 (**A**) or 2208L (**B**) treated with vehicle or Zotatifin till ethical endpoint. Scale bar, 50 μm. Right, quantification of IHC staining. Three to six representative 20X images were analyzed for each tumor. N=5 biological replicates per group. Data are presented as mean ± SEM and analyzed using two-tailed unpaired Student’s *t*-test. **C,** IHC staining of F4/80 in 2153L tumors treated with indicated therapy. Scale bar, 100 μm. **D,** Growth curves of 2153L tumors treated with indicated therapy. n=4 biological replicates for the Zotatifin+anti-Csf1r arm and n=3 for all other arms.

**Supplementary Figure 3.** Zotatifin inhibits the translation of *Sox4* and *Fgfr1* mRNAs. **A,** Immunoblotting analysis of 4T1 tumors that were treated with vehicle or Zotatifin in vivo. n = 5 biological replicates per group. **B,** Immunoblotting analysis of 2153L and E0771 cells that were treated with 40 nM Zotatifin in vitro for 1 hr or 6 hrs respectively. **C** and **D,** QPCR analysis for *Sox4* (**C**) and *Fgfr1* (**D**) in 2153L cells that were treated with different concentrations of Zotatifin for 6 hrs in vitro. **E,** Immunoblotting analysis of 2153L cells that were treated with CHX or MG132 for different time periods. **F** and **G,** QPCR analysis for *SOX4* (**F**) and *FGFR1* (**G**) in BT549 cells that were treated with 40 nM Zotatifin for different time periods. **H,** Immunoblotting analysis of 2153L cells that were treated with indicated drugs for 24 hrs. CPT, camptothecin. **I,** QPCR analysis of HAP1 cells that were treated with 40 nM Zotatifin for 6 hrs. In **C**, **D**, **F**, **G**, and **I**, data are representative of two independent experiments and are presented as mean ± SD of technical triplicates. **J,** Distribution of *Actb* and *Gapdh* mRNAs across the different fractions in polysome profiling analysis of 2153L cells that were treated with vehicle or 40 nM Zotatifin for 2 hrs. **D**ata are presented as mean ± SEM of three biological replicates. **K,** Immunoblotting analysis of 2153L cells that were treated with vehicle or Zotatifin in the presence of CHX or MG132 for 2 hrs. Data are representative of two independent experiments.

**Supplementary Figure 4.** Zotatifin induces interferon response genes. **A,** QPCR analysis of 2153L cells that were treated with different concentrations of Zotatifin for 6 hrs. **B,** QPCR analysis of 2153L cells that were treated with 40 nM Zotatifin for different time periods. In **A** and **B**, data are representative of two independent experiments and are presented as mean ± SD of technical triplicates. **C,** QPCR analysis of 2153L cells that were treated with indicated drugs for 24 hrs. Data are presented as mean ± SD of technical triplicates. **C,** QPCR analysis of paired ER+ breast cancer biopsies from pre-treatment (pre) and on Zotatifin treatment (on) patients. The mRNA levels of pre-treatment samples were set as 1 and fold changes were calculated for each paired sample. Data are analyzed using two-tailed unpaired Student’s *t*-test. n=8 patient biopsy pairs.

**Supplementary Figure 5.** Sox4 inhibition by Zotatifin contributes to Zotatifin induced interferon response genes. **A,** The mRNA levels of *Eif4a1*, *Eif4a2,* and *Ddx3x* in normal mammary glands of BALB/c mice and *Trp53*-null preclinical models. The RNA levels for each gene were averaged RNA-seq signals from 1-12 biological replicates. **B,** The Chronos dependency scores of *EIF4A1* and *EIF4A2* in CRISPR knockout screens. A lower Chronos score indicates higher essentiality. A score of 0 indicates a gene is not essential and a score of −1 is the median scores of all pan-essential genes. **C,** The RNA and protein levels of *EIF4A1*, *EIF4A2*, and *DDX3X* in non-triple-negative or triple-negative breast cancer tissues from the Clinical Proteomic Tumor Analysis Consortium (CPTAC) database. **D,** Immunoblotting analysis of 2153L cells that were transfected with siRNA for 48 hrs and left untreated or treated with 40 nM Zotatifin for the last 24 hrs. **E,** Immunoblotting analysis of BT549 cells that were transfected with siRNA for 48 hrs. * denotes a non-specific band. **F,** QPCR analysis of BT549 cells that were transfected with negative control siRNA with or without Zotatifin treatment, or *SOX4* siRNAs without Zotatifin treatment for 48 hrs. **G,** QPCR analysis of Zotatifin-induced gene fold changes in BT549 cells that were transfected with negative control siRNA or *SOX4* siRNA in the presence of vehicle or Zotatifin. In **D-G**, data are representative of two independent experiments. In **F** and **G**, data are presented as mean ± SD of technical triplicates.

**Supplementary Figure 6.** Zotatifin synergizes with chemotherapy to suppress tumor progression. **A,** Top, treatment scheme of BALB/c mice. Bottom, individual growth curves of 2153L tumors treated with indicated drugs. Each line represents a tumor and different line colors denote separate experimental batches. Data from 3 to 5 independent experimental batches are integrated. n=24 for Vehicle, n=22 for Zotatifin, n=13 for Carboplatin, and n=24 for Zotatifin+Carboplatin. **B,** Body weight changes of 2153L tumor-bearing BALB/c mice over the treatment course. n≥6 biological replicates per treatment arm. **C** and **D,** Left, representative images of IHC staining of BrdU (**C**) or cleaved Caspase 3 (**D**) in 2153L tumors that were treated with indicated drugs for 3 days. The regions outlined in box are magnified below. Scale bar, 50 μm. Right, quantification of IHC staining. Five representative 20X images were analyzed for each tumor. n = 3 biological replicates per group. Data are presented as mean ± SEM and analyzed using two-tailed unpaired Student’s *t*-test. **E,** Tumor growth curves of 2153L tumors treated with indicated drugs. n≥4 biological replicates per treatment arm. **E,** Tumor growth curves of 2153L tumors treated with indicated drugs. n=5 biological replicates per treatment arm. In **B**, **E**, and **F**, data are presented as mean ± SEM and analyzed using two-way ANOVA with Bonferroni’s multiple comparison test. **G,** Kaplan-Meier survival curves of 2153L tumor-bearing mice treated with indicated drugs. n=5 biological replicates per group. The log-rank test (two-tailed) was used to test for the significant differences of curves between groups.

**Supplementary Figure 7.** Zotatifin and carboplatin combination therapy induces interferon response genes and changes in the tumor microenvironment. **A,** QPCR analysis of 2153L cells that were treated with 40 nM Zotatifin or/ and 10 μM carboplatin for 24 hrs in vitro. **B,** UMAP plot overlaid with the expression of selected markers from mass cytometry analysis of tumor-infiltrating immune cells from all treatment groups in 2153L.

**Supplementary Figure 8.** Zotatifin and carboplatin combination therapy induces T cell infiltration to the tumor microenvironment. **A,** Representative images of IHC staining of CD4 (top) and CD8 (bottom) in 2153L tumors from BALB/c mice that were treated with indicated drugs for 11 days. Scale bar, 50 μm. **B,** Quantification of CD4 (left) and CD8 (right) IHC staining. Five to eleven representative 20X images were analyzed for each tumor. n = 3 biological replicates per group. Data are presented as mean ± SEM and analyzed using two-tailed unpaired Student’s *t*-test.

## Supplementary table legends

**Supplementary Table S1.** The proteomic alterations determined by tandem mass tag mass spectrometry in 2153L tumors treated with Zotatifin for 3 days compared with the vehicle. n=4 biological replicates per arm. Statistical significance was determined using a two-tailed unpaired moderated *t*-test.

**Supplementary Table S2.** Gene set enrichment analysis (GSEA) of pathways that are enriched in 2153L tumors treated with vehicle compared to Zotatifin using data from mass spectrometry. GSEA was performed with the MSigDB hallmarks dataset.

**Supplementary Table S3.** Gene set enrichment analysis (GSEA) of pathways that are enriched in 2153L tumors treated with Zotatifin compared to vehicle using data from mass spectrometry. GSEA was performed with the MSigDB hallmarks dataset.

**Supplementary Table S4.** Gene set enrichment analysis (GSEA) of pathways that are enriched in 2153L tumors treated with Zotatifin+carboplatin compared to Zotatifin monotherapy using data from mass spectrometry. GSEA was performed with the MSigDB hallmarks dataset.

**Supplementary Table S5.** Gene set enrichment analysis (GSEA) of pathways that are enriched in 2153L tumors treated with Zotatifin+carboplatin compared to carboplatin monotherapy using data from mass spectrometry. GSEA was performed with the MSigDB hallmarks dataset.

**Supplementary Table S6.** Sequences of qPCR primers.

