## Supplementary Figures for "Targeting EIF4A triggers an interferon response to synergize with chemotherapy and suppress triple-negative breast cancer"

### Supplementary Figure 1

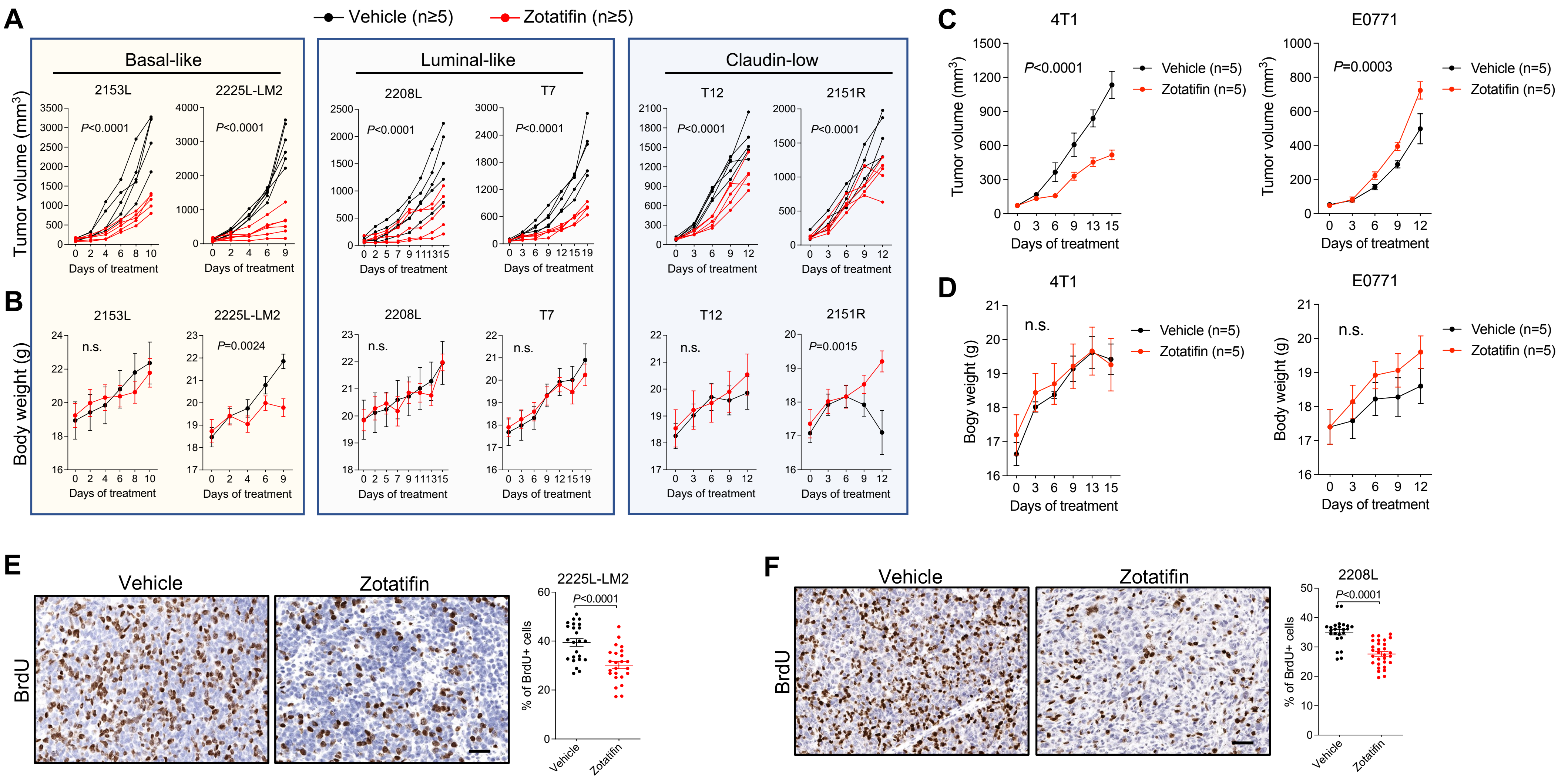

Supplementary Figure 2

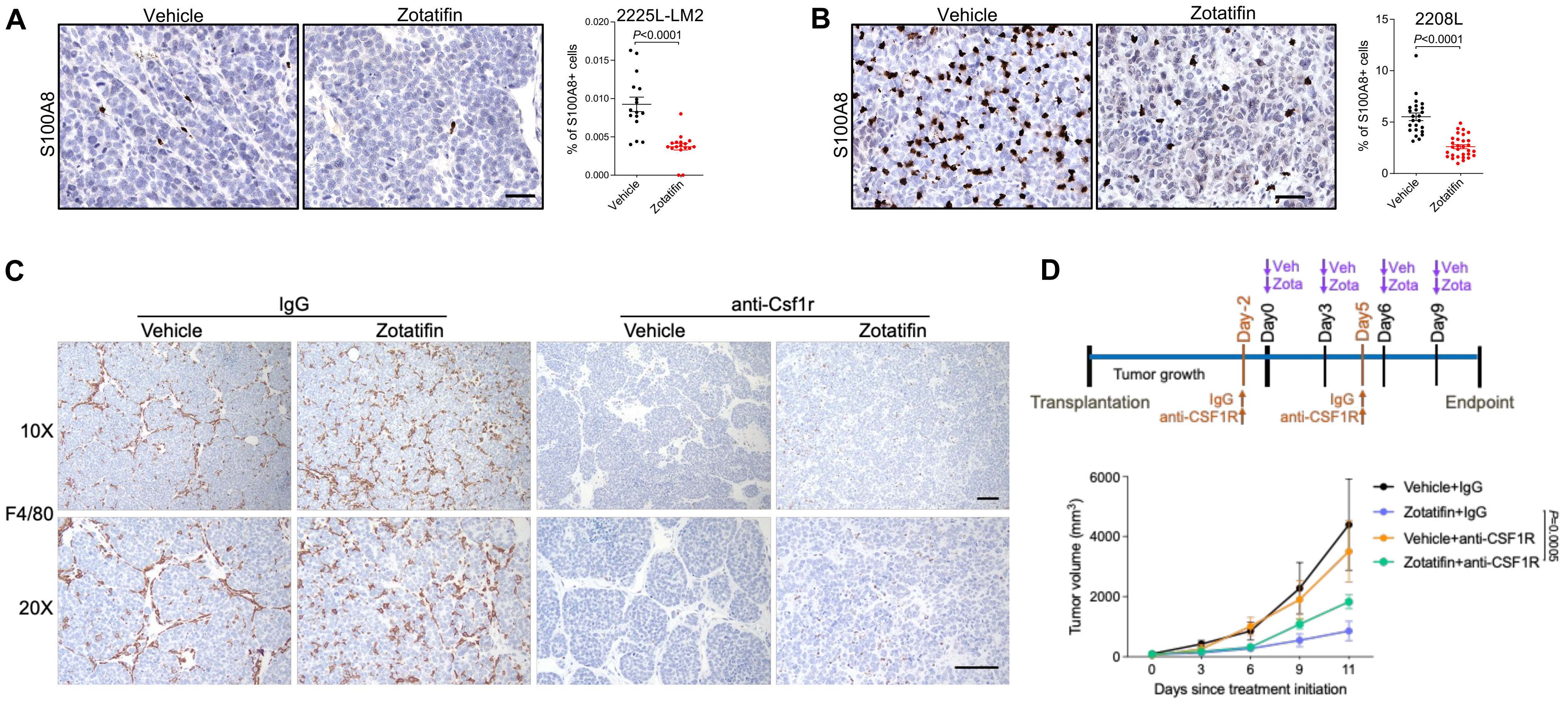

### Supplementary Figure 3

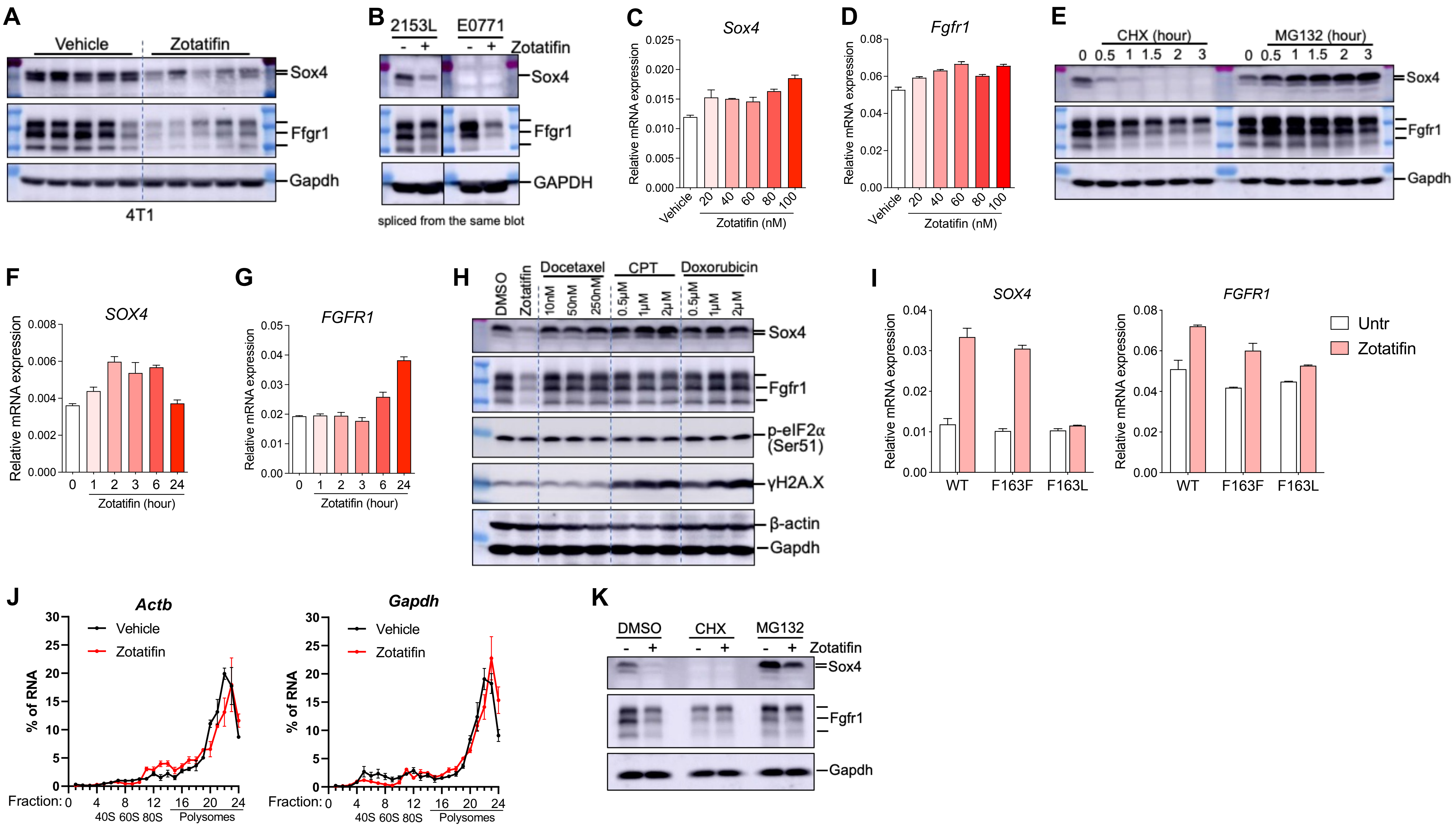

### Supplementary Figure 4

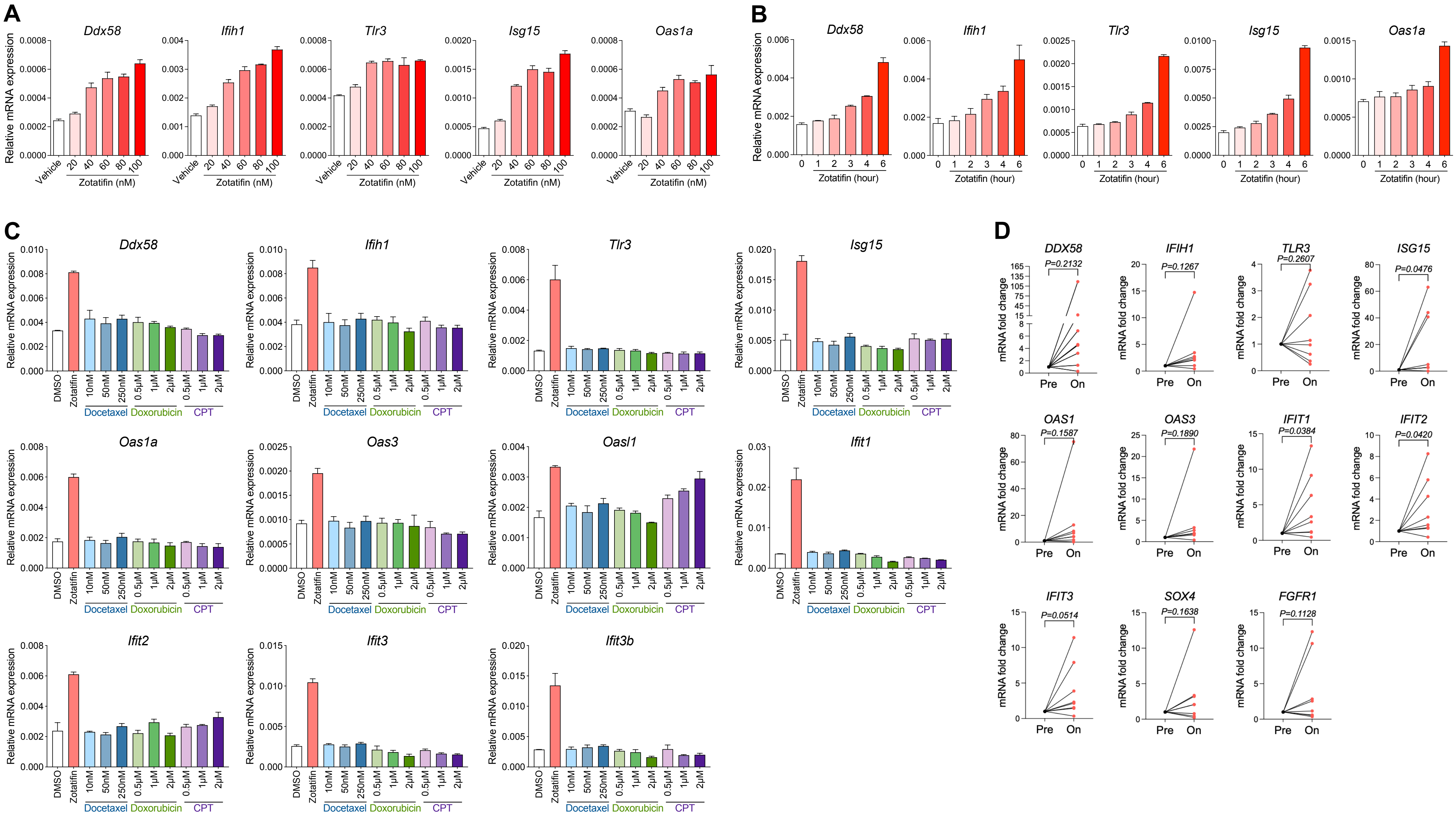

### Supplementary Figure 5

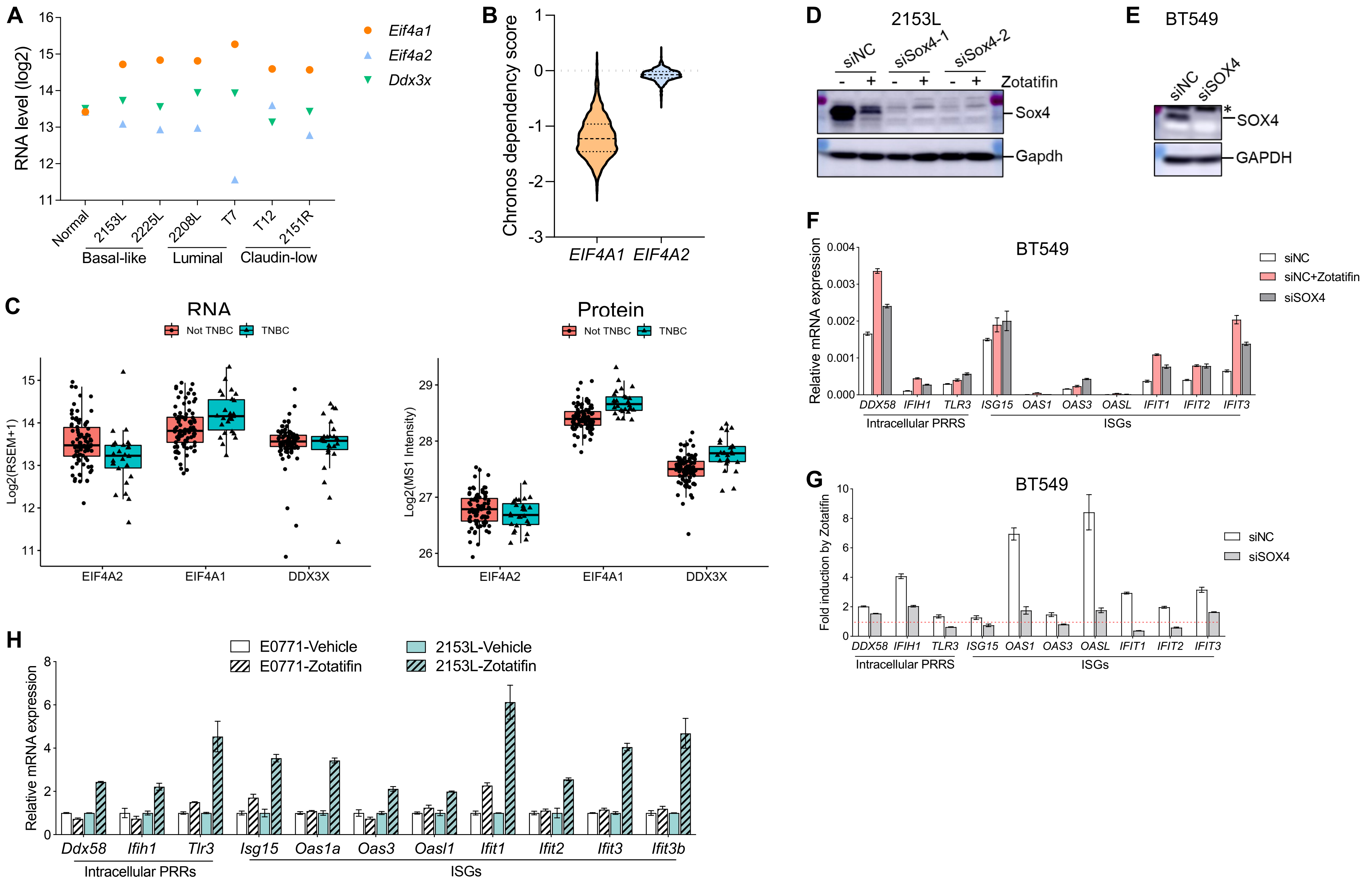

### Supplementary Figure 6

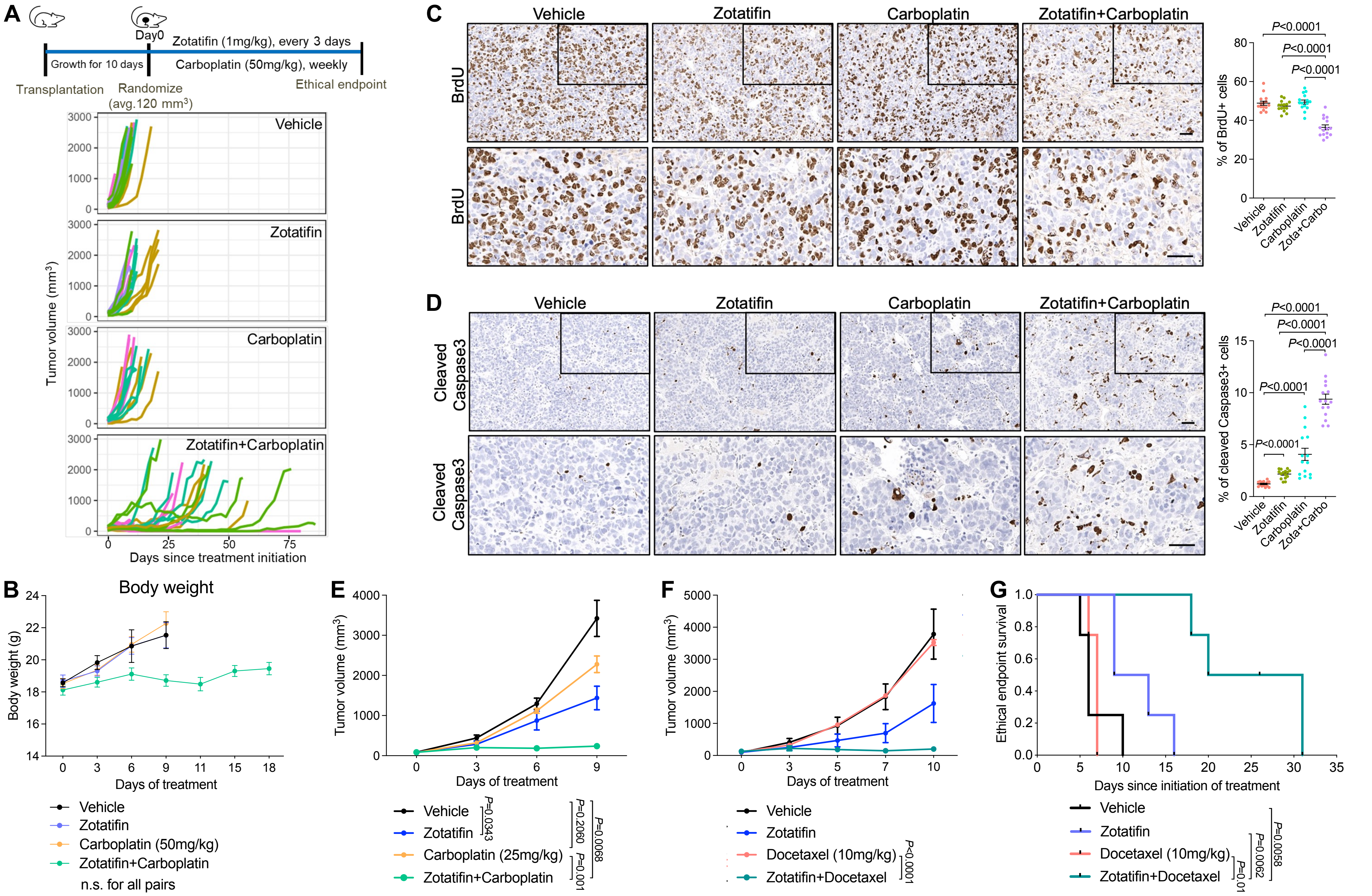

Supplementary Figure 7

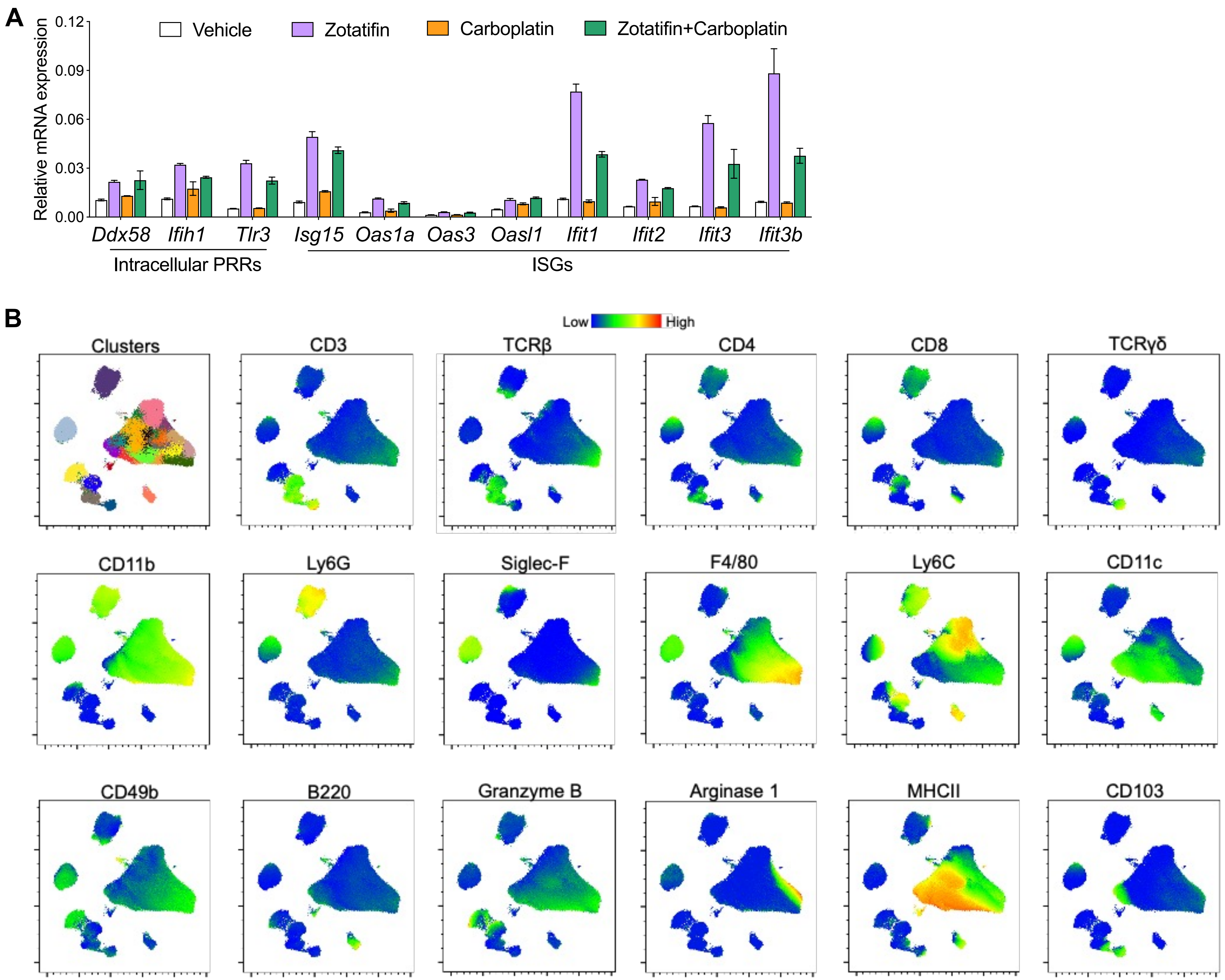

Supplementary Figure 8

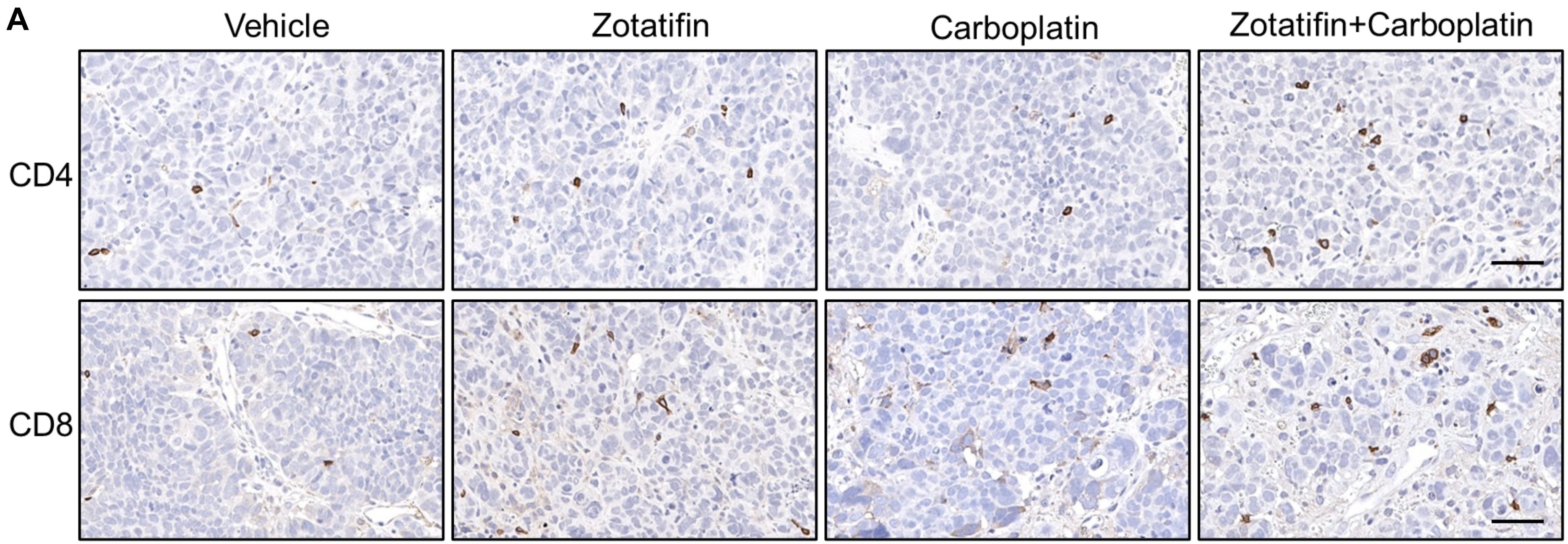


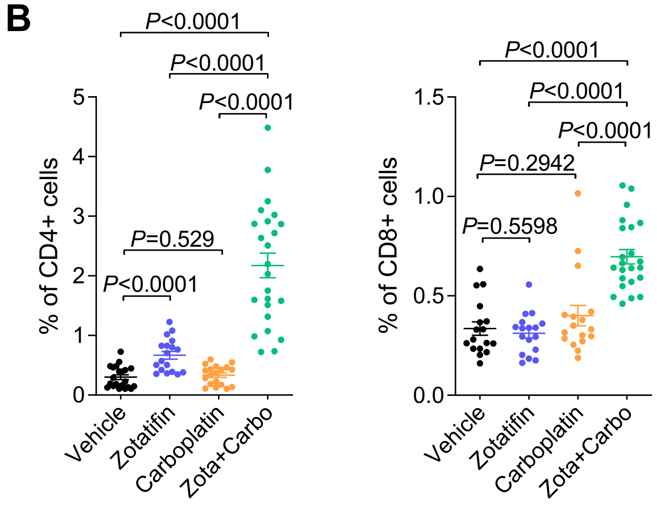
